# Terminal heterocyst differentiation in the *Anabaena patA* mutant as a result of post-transcriptional modifications and molecular leakage

**DOI:** 10.1101/2021.12.20.473559

**Authors:** Pau Casanova-Ferrer, Saúl Ares, Javier Muñoz-García

## Abstract

The *Anabaena* genus is a model organism of filamentous cyanobacteria whose vegetative cells can differentiate under nitrogen-limited conditions into a type of cell called heterocyst. These heterocysts lose the possibility to divide and are necessary for the colony because they can fix and share environmental nitrogen. In order to distribute the nitrogen efficiently, heterocysts are arranged to form a quasi-regular pattern whose features are maintained as the filament grows. Recent efforts have allowed advances in the understanding of the interactions and genetic mechanisms underlying this dynamic pattern. However, the main role of the *patA* and *hetF* genes are yet to be clarified; in particular, the *patA* mutant forms heterocysts almost exclusively in the terminal cells of the filament. In this work, we investigate the function of these genes and provide a theoretical model that explains how they interact within the broader genetic network, reproducing their knock-out phenotypes in several genetic backgrounds, including a nearly uniform concentration of HetR along the filament for the *patA* mutant. Our results suggest a role of *hetF* and *patA* in a post-transcriptional modification of HetR which is essential for its regulatory function. In addition, the existence of molecular leakage out of the filament in its boundary cells is enough to explain the preferential appearance of terminal heterocysts, without any need for a distinct regulatory pathway.

**Author summary:** Understanding multicellular pattern formation is key for the study of both natural and synthetic developmental processes. Arguably one of the simplest model systems for this is the filamentous cyanobacterium *Anabaena*, that in conditions of nitrogen deprivation undergoes a dynamical differentiation process that differentiates roughly one in every ten cells into nitrogen-fixing heterocysts, in a quasi-regular pattern that is maintained as the filament keeps growing. One of the most characteristic mutations affecting this process forms heterocysts mostly constrained to the terminal cells of the filament. We have used experimental observations to propose a mathematical model of heterocyst differentiation able to reproduce this striking phenotype. The model extends our understanding of the regulations in this pattern-forming system and makes several predictions on molecular interactions. Importantly, a key aspect is the boundary condition at the filament’s ends: inhibitors of differentiation should be able to leak out of the filament, or otherwise the terminal cells would not differentiate. This highlights, in a very clear example, the importance of considering physical constraints in developmental processes.

## Introduction

The biology of cyanobacteria has been the subject of intensive work during the last decades. A great number of studies have focused on a specific type of cyanobacteria that forms colonies consisting of one-dimensional filaments, the strain PCC 7120 of the genus Anabaena, to such an extent that this has become a model organism in the field [1–3]. Under nitrogen-rich conditions, these filaments are composed only of vegetative cells carrying photosynthesis. However, as a response to different environmental stresses, vegetative cells can differentiate into specialized cell types that can fix atmospheric nitrogen. These individual conversions allow the survival of the colony and represent a paradigmatic example of a prokaryotic life form with differentiated cell types. Furthermore, bacteria and archaea are the only organisms capable of fixing atmospheric nitrogen, which makes them crucial for the viability of all living beings on Earth. The process used during nitrogen fixation in cyanobacteria is catalyzed by nitrogenase, and this enzyme is easily degraded by oxygen. In order to avoid this degradation, some cyanobacteria have developed a mechanism to protect nitrogenase from the oxygen produced by vegetative cells. This mechanism is the specialization of some cells into a form denominated heterocysts which accumulates nitrogenase and does not carry out photosynthesis. When external nitrogen sources are scarce, heterocysts appear in a quasi-regular pattern, with intervals of roughly ten vegetative cells between heterocysts. Fixed nitrogen produced by the heterocysts reaches the vegetative cells and sustains their growth. Reciprocally, nutrients produced by photosynthesis in vegetative cells are also shared to maintain the production of nitrogenous compounds in heterocysts, which require high energy consumption [1,4]. Upon differentiation, heterocysts lose the possibility to undergo cell division, but vegetative cells continue dividing, producing filament growth and increasing the distance between consecutive heterocysts. Due to this, new heterocysts differentiate in the middle of the intervals between previously existing heterocysts in order to not diminish the supply of nitrogen to distant vegetative cells. In this way, the dynamic process of differentiation allows the overall pattern of heterocyst location to conserve its properties over time.

There are a large number of processes involved in the regulation of heterocyst pattern formation and its maintenance. In addition to nitrogen levels and other environmental aspects, many genes and transcription factors play a role [5]. It has been shown that, when nitrogen stress is perceived, the transcription regulator *ntcA* is important to trigger heterocyst differentiation [6,7], by directly or indirectly controlling the expression of several genes including *hetR* [8]. The expressions of *ntcA* and *hetR* are mutually dependent, although the latter seems to be sufficient for heterocyst development [9]. Thus, positive auto-regulation of *hetR* is required for differentiation and is particularly significant in developing heterocysts [10,11]. *hetR* expression is the main positive regulatory factor in heterocyst development [9,10] and this gene is epistatic to the others involved in heterocyst differentiation [12].

The gene *patS* is a negative regulator of *hetR*, suppressing differentiation when overexpressed and inducing multiple contiguous heterocysts, the so-called Mch phenotype, when deleted [13,14]. The expression of *patS* produces a short peptide PatS, predicted to be formed by 13 or 17 amino acids, which contains a carboxyl-terminal that prevents DNA binding activity of HetR [15, 16] and inhibits differentiation when added to culture medium [13]. The expression of *patS* in small groups of cells was shown to diminish the levels of HetR in adjacent cells [17], suggesting that a PatS-dependent signal can be trafficked along the filament [18]. Recent studies have also proven a redundancy in the inhibitory signal of *patS* through the gene *patX* [19,20]. Despite acting in an analogous way to *patS, patX* seems to play a secondary role complementing the main signal produced by *patS*. As shown in [20], the Δ*patX* single mutant does not present an altered phenotype while the Δ*patX* Δ*patS* double mutant displays a much higher percentage of heterocysts than the Δ*patS* single mutant.

Although lack of *patS* expression initially produces a pattern with groups of contiguous heterocysts and short intervals of vegetative cells between those clusters, this pattern tends to recover the characteristics of a wild-type-like pattern later on [14]. This suggests the presence of additional patterning signals operating long after nitrogen deprivation. The most relevant player that leads to this late inhibitory effect is the *hetN* gene, expressed only in heterocysts. Similarly to *patS*, the product codified by *hetN* also contains an ERGSGR motif, raising the possibility that an ERGSGR-containing peptide derived from the full protein goes from cell to cell [21,22]. However, in contrast to the *patS* mutant phenotype, the *hetN* mutant phenotype has a heterocyst pattern similar to the wild-type at the initial stages of nitrogen depletion and a multiple contiguous heterocyst phenotype after 48 hours [23]. This indicates that *hetN* expression is activated later than that of *patS*. Additional proof of the inhibitory function of *patS* and *hetN* is that, when both genes are suppressed, almost all cells eventually differentiate, causing lethal levels of heterocysts [24].

Experimental results, such as the previously described and more recent ones with transcriptional information [25], have allowed advancing in the understanding of the mechanisms and interactions between *hetR, patS*, and *hetN* that give rise to the appearance and maintenance of heterocyst patterns. However, in addition to these, other transcription factors such as *patA* and *hetF* have been shown to play an important role at the early steps of differentiation, regulating the transcriptional activity of *hetR* [26]. All this complex network of interactions, where other heterocyst-related genes, such as *hetC, hetP, hetL, patN*, and *hetZ*, also play a role, has made the complete understanding of heterocyst differentiation a challenge during the last two decades.

In this work, we investigate the function of *patA* and *hetF* together with their interactions with the main genes responsible for heterocyst pattern formation. In the next section we review the main experimental results regarding these genes. Based on these findings we propose a theory combining genetic, metabolic, and morphological aspects. The proposed mathematical model reproduces the diverse experimental phenotypes and explains the main function of both *patA* and *hetF* in the gene-regulatory network of heterocyst differentiation.

### Experimental evidence about *patA*

The gene *patA* and its product, together with its mutant phenotype, were first described by Liang, Scappino, and Haselkorn [27]. The most striking feature of the mutant is that it forms heterocysts almost exclusively at the terminal cells of the filament. In this seminal work, it was described that the *patA* gene product contains a carboxyl-terminus similar in sequence to CheY (a protein subject to phosphorylation that promotes rotation of bacterial flagella in *Escherichia coli* [28]) and an amino-terminus with the so-called PATAN domain (predicted to participate in protein-protein interactions [29]). *patA* is more abundantly transcribed in nitrogen starvation conditions [27], with levels of expression on proheterocysts slightly larger than in vegetative cells [30]. In regards to the observable phenotype, *patA* mutation suppresses heterocyst differentiation primarily in the intercalary cells, in most cases forming only a pair of heterocysts at both ends of the filament. The fact that *patA* seems to be required for the differentiation of intercalary heterocysts but not for terminal heterocysts have made some authors think that a different differentiation process in which *patA* is not involved could occur depending on cell position [12].

A deeper understanding of the function of *patA* can be obtained by noticing that its overexpression gives rise to changes in the morphology of vegetative cells and a decrease in the average number of vegetative cells between heterocysts. In [30] it is shown that a strain overexpressing *patA* presents 15.5% of heterocysts 48 hours after nitrogen depletion, in contrast to the smaller percentage of 7.6% obtained for the wild-type in the same conditions. This information suggests that PatA moderately promotes heterocyst differentiation, reducing the distance between consecutive heterocysts. In this work, it is also stated that *patA* overexpression produces an enlargement of the vegetative cells and disruptions of the division plane. Since *patA* seems to be located predominantly at sites of cell division, it is suggested that *patA* might modify the communication conduits between cells, thus altering the exchange of metabolites and regulatory molecules.

Regarding the connection between *patA* and the master regulator in heterocyst differentiation, *hetR*, multiple contiguous heterocysts appear when *hetR* is overexpressed [27]. However, this differentiation is mostly suppressed in the *patA* mutant, for which the same *patA* phenotype with only terminal heterocysts is obtained even under nitrogen starvation. A similar result is reported in [31], where a copper inducible *petE::hetR* fusion is used to check that addition of Cu^2+^ does not induce the formation of heterocysts in a *patA* mutant. These results indicate that a functional *patA* is required for the normal function of *hetR*. However, as we will see below, this interaction is not direct. In [31] the authors conclude that *patA* does not directly increase *hetR* transcription and suggest that this gene is probably involved in other post-transcriptional steps.

Conversely, more recent experiments have shown that a functional *hetR* gene is required for the induction of expression of *patA*. Using PatA-GFP and *β*-galactosidase activity, in [30] it was observed that the *patA* transcription is greatly reduced in strains for which the expression of *hetR* is blocked. Therefore, the activation of *patA* expression seems to be directly upregulated by HetR. Actually, in [32] evidence was found that HetR binds directly to the promoter of *patA*. Also, in [33], it is claimed that *patA* activation is controlled by NtcA. Since the expressions of both *hetR* and *ntcA* are higher after nitrogen step-down and mutually positive regulated [4], this positive interaction would directly explain the higher abundance of *patA* observed in nitrogen starvation conditions.

Surprisingly, even though rare intercalary heterocysts are formed in strains lacking the *patA* gene, high levels of HetR are measured 18 hours after removing combined nitrogen [26]. These levels are higher than in the wild-type, as measured by western blot analysis using a polyclonal antibody directed against HetR. The increase in these levels in the *patA* mutant is also confirmed in [34] using HetR-GFP translational fusion controlled by a copper-inducible *hetR* promoter. These results seem to contradict the idea that high levels of HetR necessarily imply the formation of more frequent heterocysts. The HetR-GFP fluorescence gradients in this work also show that there is a post-transcriptional *hetR* regulation through the ERGSGR hexapeptide that seems to increase HetR degradation.

More insights into the functional relationship between *patA* and other genes involved in heterocyst differentiation are presented in [12]. This work studies the connections between *patA* and *hetR, patS*, and *hetN*, analyzing the single, double, and triple mutant phenotypes. For those double mutants in which *hetR* was knocked out, they observe the same phenotype without heterocyst differentiation as in the single *hetR* mutant, confirming that *hetR* is epistatic to *patA, patS* and *hetN*. For the *patA* mutant background with *hetN* inactivated, they obtain a phenotype indistinguishable from the *patA* mutant with single terminal heterocysts at 24 hours post-induction. However, an increasing number of contiguous heterocysts are formed mostly at the ends of the filament after that time.

In the case of the Δ*patA*Δ*patS* double mutant, its phenotype is identical to the Δ*patS* single mutant during short times after induction. The double mutant presents a similar number of heterocysts and an equivalent length of vegetative intervals but without the typical increase of HetR concentration of the rest of studied Δ*patA* mutants. This seems to imply that a functional *patS* gene is required to obtain a *patA* phenotype. However, after 48 hours the average distance between heterocysts for the Δ*patA*Δ*patS* double mutant is larger than in the Δ*patS* single mutant and the wild-type. This double mutant also presents a phenotype resembling a *patA* mutant in the intervals between primal heterocysts [12]. In the case of filaments overexpressing both *hetR* and *patS*, the authors found that they do not present heterocysts in a wild-type genetic background. Hence, it seems that the overexpression of *patS* is epistatic to the overexpression of *hetR*.

Based on the previous results, it is suggested that PatA might reduce the efficiency of the inhibitory function of both PatS and HetN. This effect could be achieved in two general ways. The first one would be that PatA interacts with PatS and HetN to reduce its inhibitory potential through a post-transcriptional modification that could be forced degradation, a conformation change, or sequestration. On the other hand, PatA could also interact with HetR to protect it by reducing its sensitivity to inhibition. Nevertheless, given that the Δ*patA*Δ*patS* and the Δ*patA*Δ*patS*Δ*hetN* mutants do not present the same phenotype of the Δ*patS* and the Δ*patS*Δ*hetN* mutants, PatA must have another effect besides the protection of HetR to the inhibition through PatS and HetN. The Δ*patA*-like phenotype obtained for an isolated allele of *hetR* made the authors in [12] suggest that *patA* might also promote differentiation independently from its effects on *patS* and *hetN*.

### Experimental evidence about *hetF*

The first identification of the gene *hetF* was made on another species of cyanobacteria, *Nostoc punctiforme*, by Wong and Meeks [35]. This work shows that the Δ*hetF* mutant does not form heterocysts in nitrogen starvation conditions. It is also stated that overexpression of *hetF* increases the frequency of heterocysts in the wild-type and Δ*hetF* mutant under nitrogen deprivation. Moreover, the *hetR* mutant cannot be rescued with overexpression of *hetF*. Additionally, *hetF* overexpression produces changes over the filament morphology, which becomes irregular due to cell septation out of the normal plane of division. Lastly, they found that *hetF* is always transcribed constitutively at a low level and that HetR accumulates non-specifically in the Δ*hetF* mutant.

A posterior study by Risser and Callahan [26] shows that the deletion of *hetF* in *Anabaena PCC 7120* produces enlarged vegetative cells (with a morphology similar to *patA* overexpression) that do not differentiate into heterocysts even after several days on nitrogen starvation. When *hetF* is overexpressed, vegetative cells become significantly smaller than those in the wild-type, and multiple contiguous heterocysts (the Mch phenotype) is induced 24h after nitrogen step-down. This work also studied the interaction between *patA* and *hetF*. Their deletion mutants (and the double mutant) present similar high levels of HetR. However, the addition of an ectopic functional HetF reverts the phenotype of all these mutants to the wild-type. Furthermore, the regulatory effect of those genes on *hetR* seems to be post-transcriptional. Both *patA* and *hetR* are necessary for the aberrant cell morphology of the Δ*hetF* mutant and the addition of extra copies of *hetF* can functionally bypass the deletion of *patA* without requiring a direct interaction of PatA with *hetF*. Finally *hetR* self-regulation and *patS* upregulation through HetR depend on *hetF*. These results lead the authors of [26] to suggest the existence of an activation process of HetR controlled by *hetF* which induces the *hetR* regulatory function.

## Materials and methods

### Mathematical Model

A number of works have presented mathematical models of heterocyst differentiation [36–49]. Here we have used the interactions explained in the gene regulatory network section and depicted in Fig 1 to formulate our model schematically presented in Fig 2. Based on these interactions we formulated a set of differential equations for the evolution of the species involved in heterocyst differentiation. Assuming that protein interactions are much faster than the production of those proteins, equilibrium states were considered for the reactions with a shorter timescale. Applying these simplifications we get a more manageable mathematical model to describe the temporal evolution of the concentration of the main protein monomers. Thus, the temporal evolution equation for the main species are

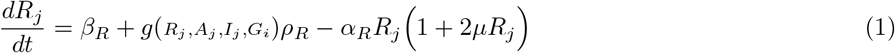

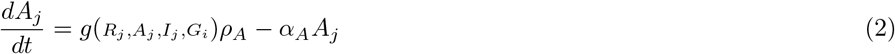

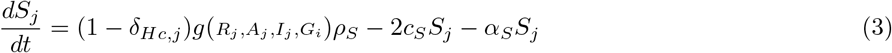

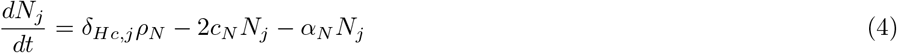

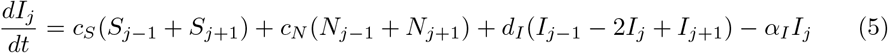

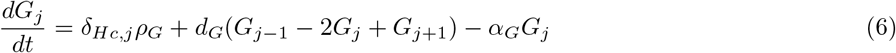

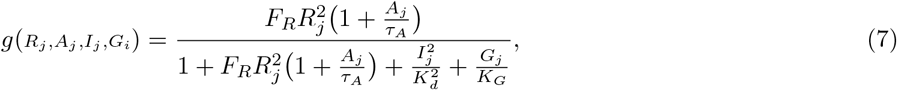

where the subindex *j* indicates that the variable refers to the cell number *j* in the filament (being then *j* – 1 and *j* + 1 its neighboring cells) and *t* denotes time. The concentration of the protein monomers are represented by *R_j_*, *A_j_*, *S_j_*, and *N_j_*, which stand for the concentration of HetR, PatA, the addition of both PatS and PatX, and HetN, respectively, in the cell *j*. As explained in the gene regulatory network section, we assume for both inhibitory genes (PatS and HetN) the same functional form, the hexapeptide ERGSGR, represented by *I_j_* in our model. Finally, *G_j_* represents the concentration of fixed nitrogen in cell *j*.

**Fig 1.**
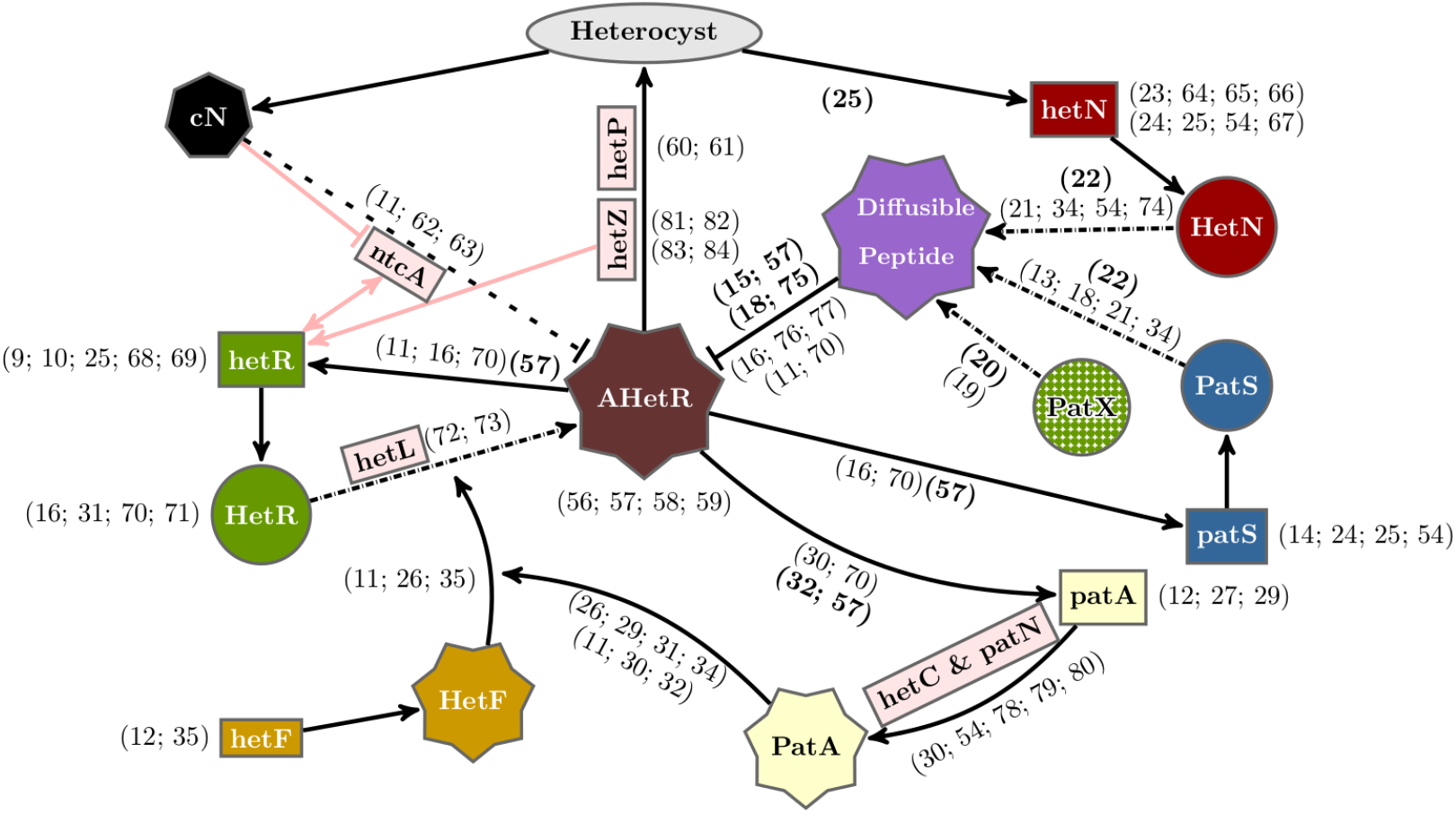
Gene regulatory network of heterocyst differentiation. The main elements involved and their interactions for heterocyst differentiation are depicted schematically together with other relevant genes (light red rectangles) not included in our model. Rectangles, circles, and polyhedric forms represent genes, inactive proteins, and active products, respectively. AHetR stands for the active form of HetR. The ellipse represents differentiation into a heterocyst. Arrows with solid lines represent interactions between elements. Arrows with dashed-dotted lines represent post-transcriptional changes. The dashed line represents a simplified interaction of nitrogen sensing through *ntcA*. References justifying these interactions are incorporated. Regular and bold formatted references indicate phenotypically inferred and experimentally observed interactions, respectively.

**Fig 2.**
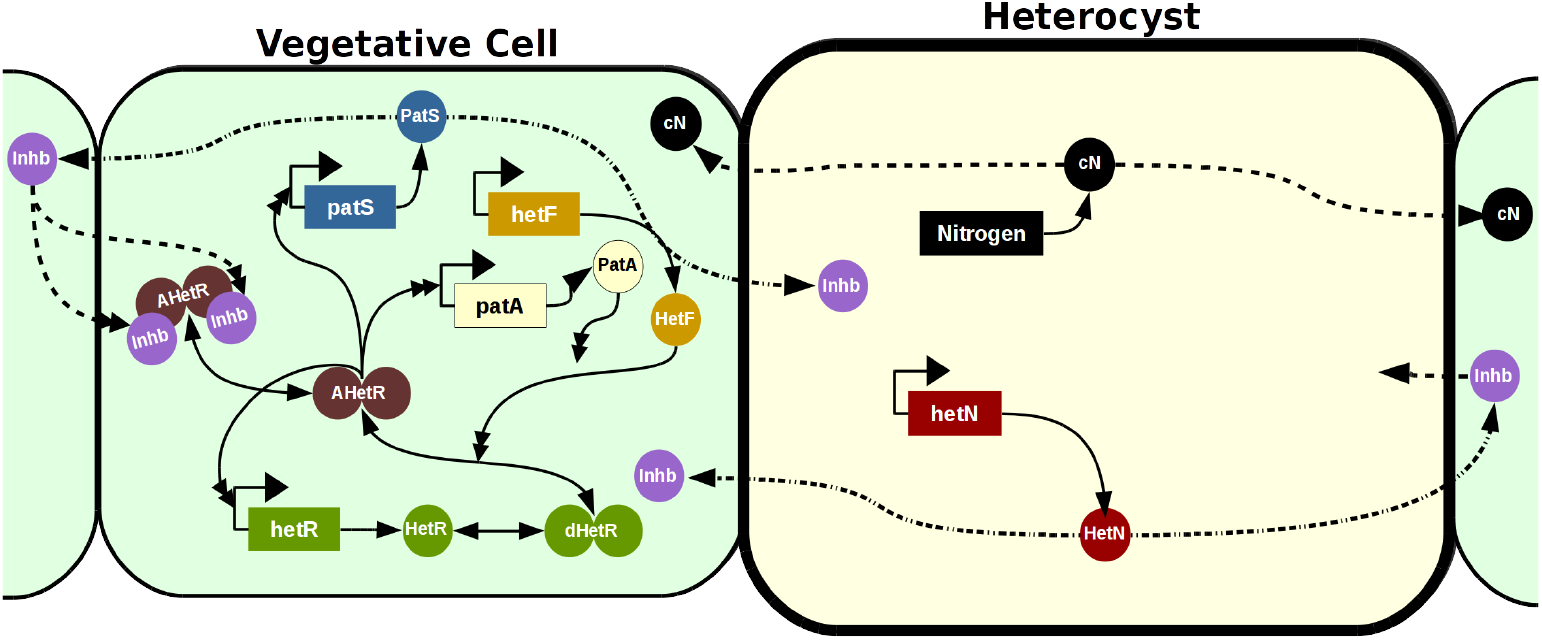
Mechanistic Model. The vegetative cells are represented with a soft green background while the heterocyst has a soft yellow background and a thicker cell wall. Genes are represented in rectangles and proteic elements with circles. The dimers are represented with two attached circles and can be inactivated (in green), activated (in brown), and activated and inhibited (in brown with two attached purple inhibitors). Solid lines represent production (with only one simple arrowhead), transformations (with a simple arrowhead in both ends), and interactions (with a double arrowhead). Dashed lines represent inter-cellular traffic and dashed-doted lines represent a transformation when exported to a neighboring cell.

Assuming that HetF is produced at a constant basal rate and degraded linearly, we have simplified the model considering an equilibrium concentration of HetF (*F_eq_*) following

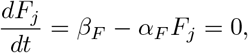

from where we get a constant concentration of HetF at any cell: *F_eq_* = *β_F_/α_F_*.

For simplicity, we have only considered basal production of HetR monomers and HetF proteins with rates *β_R_* and *β_F_* respectively. The maximum regulated production rates are represented by *ρ_R_, ρ_A_, ρ_S_*, *ρ_N_*, and *ρ_G_*, for HetR, PatA, both PatS and PatX, HetN, and fixed nitrogen, respectively. The linear degradation rates are *α_R_, α_F_*, *α_A_, α_S_*, *α_N_, α_I_*, and *α_G_*, and the active transport rates or diffusion between adjacent cells are *c_S_, c_N_*, *d_I_*, and *d_G_*. We assume the border cells at the filament’s ends leak both the inhibitor and the fixated nitrogen to the exterior at a lower rate than the communication between neighboring cells. We have modeled this multiplying the rates *d_I_*, *d_G_* by a factor *d_border_*. The exact value of this factor does not have a qualitative effect on the model but can affect the number of terminal heterocysts in some conditions.

The *μ* parameter refers to the nonlinear degradation of HetR mediated through its dimerization and could be further expressed as a function of the rates of binding (*k_b_*) and unbinding (*k_u_*) of monomers to form dimers and the degradation rates for both monomers (*α_R_*) and dimers (*α_d_*) of HetR as

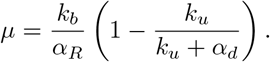

The factor *δ_Hc,j_* specifies whether the cell is vegetative or a heterocyst: its value is 1 if cell *j* is a heterocyst, 0 if it is vegetative. This value is used as a switch for the production of both HetN and fixed nitrogen which are only produced in heterocysts.

We can simplify the system assuming that the promoter regulation through HetR is equivalent for *hetR, patA* and *patS*. This regulation is modeled using the factor *g*(*R_j_, A_j_, I_j_, G_i_*). Equation 7 represents the equilibrium state of the processes of dimerization, activation, and inhibition of HetR. To obtain this expression we have considered the following equilibrium constants: *K_R_* for the dimerization of HetR, *K_F_* for the activation of the HetR dimers by HetF, *τ_A_* for the activation mediated through PatA, and *K_d_* and *K_G_* for the inhibition through the hexapeptide and the fixed nitrogen respectively. Assuming an equilibrium concentration of HetF, the expression for this regulatory term is reduced to

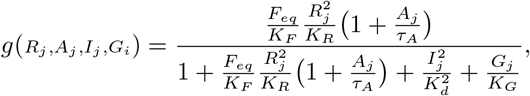

where we have defined

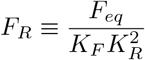

in order to obtain the simplified version in Eq 7.

### Model Implementation

We have implemented a code in an object-oriented platform to model both the biochemical interactions which give rise to heterocyst differentiation and filament growth. Each cell of the filament has its own variables representing the cellular size and concentration of considered species. The dynamical equations for the concentrations of ERGSGR inhibitor and fixed nitrogen in each cell are coupled with its adjacent neighbors. The resulting set of equations that controls the filament evolution is the noisy extension of the deterministic system Eqs 1–7. This system of equations has been expanded to the Langevin dynamics in the Itô interpretation [50]. This expansion adds an stochastic term of the form 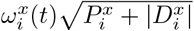 for each cell *i* and species *x*, where 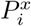 and 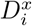 are the sum of production (synthesis) terms and the sum of degradation terms respectively and 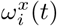 is an uncorrelated Gaussian white noise [51]. This noise has zero mean and variance 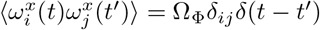 and models the intrinsic fluctuations in the genetic dynamics.

Vegetative cell growth was modeled by a stochastic differential equation for each cell:

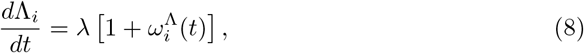

where Λ_*i*_ is the size of cell *i*, λ is a constant growth rate, and 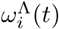 is an uncorrelated Gaussian white noise with zero mean and variance 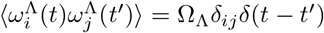, which models the intrinsic fluctuations in the growth process. Starting from an initial size, each cell evolves following Eq 8 up to a maximum size 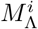, which is a noisy value drawn for each cell from a Gaussian distribution of mean *M*_Λ_ and variance 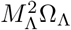. We have used Ω_Λ_ to parametrize this variance for simplicity, to avoid having too many parameters describing noisy magnitudes.

When this size 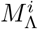 is reached, the vegetative cell divides, producing two new vegetative cells with one-half of its current size and identical protein concentrations. Heterocysts follow the same growth, but once they have reached their maximum size, 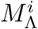, they do not divide and stop growing.

To differentiate into a heterocyst, a vegetative cell has to accumulate up a certain level of HetR. This has been implemented with an integration of the value of HetR concentration over time for each vegetative cell, once the value of *R_j_* is above a threshold, 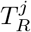.

This threshold is cell-specific, being drawn from a Gaussian distribution with mean *T_R_* and variance 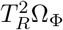. If at any point the value of *R_j_* drops below 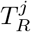, the integral is reset to zero. Otherwise, if the integral ever reaches a value 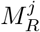, also a cell-specific Gaussian distributed parameter with mean *M_R_* and variance 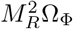, the vegetative cell differentiates into a heterocyst. Given the integrative nature of this process, we have set a minimum time *T_min_* necessary to avoid unrealistic sudden differentiation due to spikes in HetR production and properly reflect the extensive biological changes required to obtain a mature functional heterocyst. For simplicity, we have used Ω_Φ_ to parametrize variability of *T_R_* and *M_R_*, again to avoid defining too many noise-related parameters.

The parameter values employed in simulation results shown in this work can be found in Table.S1. The value of the affinity of ERGSGR to HetR, represented by *K_d_*, was taken from [15]. Using the value of 4 *μ*m for the mean maximum cell size, the value considered for the cellular growth rate, represented by λ, was chosen to agree with filament growth data in [52]. In order to obtain the parameter values that best fit the experimental data, a custom simulated annealing algorithm [53] was employed for those indicated over the double line in Table.S1, employing as initial values for the optimization, those in the model of [47] when an equivalent parameter exists.

To simulate loss-of-function conditions we have considered the production rates equal to zero, except for of the *patS* loss-of-function, where we have reduced the production rate of *patS* by 90%. The remaining 10% represents the redundant effect still present through the expression of *patX* [19,20].

### Sensitivity analysis

To assess the robustness of our results, we have performed a sensitivity analysis following the approach in [47]. We calculate the sensitivity of the model to a given parameter *X* by evaluating the observable *Y* at two points, the wild-type value *X*_0_ and the perturbation *X* = *X*_0_ + Δ*X*. Using the resulting change in the observable, Δ*Y*, we calculate the sensitivity *S_YX_* of the observable to the parameter as:

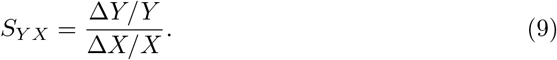

In Fig S1.Fig we show the sensitivity of the mean distance between contiguous heterocysts to changes of 10% in the parameters. The results are qualitatively similar to our previous work [47], however, we find that the extension of the model has made it even more robust to variations in individual parameters. The largest sensitivity is found when modifying HetR production and degradation, followed by inhibitor degradation, the strength of the inhibition, and PatS production. For all other parameters, the relative changes in mean interval length are much smaller than the relative change in the parameter.

### Boundary condition mechanism for the *patA* phenotype

A simple physical analogy of the importance of the boundary condition to understand the *patA* phenotype can be made using the following continuous reaction-diffusion model defined on a filament of length *L* where the position is denoted by the coordinate *x* ∈ [0, *L*]:

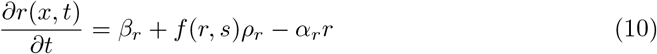

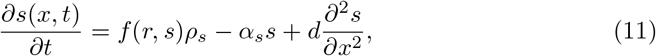

where *r* is the concentration of a non-diffusing activator (HetR), *s* the concentration of a diffusible inhibitor, *β_r_* a basal production for the activator (*β_r_* > 0), *f*(*r, s*) a smooth regulatory function, monotonically increasing in *r* and decreasing in *s*, and the other symbols are parameters. These equations need to be supplemented with boundary conditions for *s*. If the inhibitor cannot diffuse across the filament’s boundaries, the condition is:

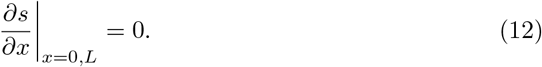

With this condition, we assume that the system is such that a parameter regime exists where there is a stable homogeneous solution. However, if leakage of the inhibitor through the boundaries is possible, the relevant boundary condition is:

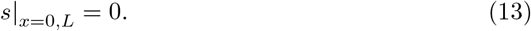

With this condition and under the same parameter regime, there would be a gradient of inhibitor decreasing from the center to the boundaries of the filament, which in turn would produce a profile of activator with maxima in the boundaries, making them the favored location for heterocyst differentiation. This simple analogy explains the physical mechanism behind the boundary-induced pattern observed in the *patA* mutant and agrees qualitatively with the observation in the discrete model depicted in Fig 6.

### Statistics

In order to compare the results obtained from our numerical simulations with the experimental data from [12, 54] we have replicated the statistical analysis of these works. In both cases, all the data is aggregated for each experiment and then the averages and standard deviation are calculated between the experimental replicates. Thus, a certain amount of filaments have been aggregated in batches and then averaged to obtain the standard deviation. On the experimental side, in [54], for each strain, 300 cells, or 100 intervals, were counted in three or four independent experiments. In [12], the number of contiguous heterocysts at the ends of 50 filaments is averaged in three experiments. Alternatively, our data was obtained from 15 batches of 10 simulations for filaments with an initial size of 30 cells (consistent with experiments in [55]) which grow to have around 50, 100, 200 cells at 24h, 48h, and 72h respectively. Therefore, we present a bigger data sample than the one considered in [54] and of the same order of magnitude as the one in [12].

## Results and Discussion

### Gene regulatory network

Using the information presented in the Introduction we propose a regulatory network, depicted in Fig 1, that includes the main genes involved in heterocyst differentiation. This genetic network modifies and expands a previous minimal model for the interaction of *hetR, patS*, and *hetN* [47].

This proposal includes the novel gene *patX*. This gene has recently been described as a redundant gene for the inhibitory mechanism of *patS* [19,20]. It has been also shown that, while its deletion in the wild-type background does not alter considerably the phenotype, the double mutant Δ*patSpatX* is lethal with an almost complete Mch phenotype [19, 20]. To take this into account in a simple way, we consider that the variable in our model for PatS represents the combined effects of PatS and PatX. The knock-out of *patS* will be modeled reducing 90% the value of the production rate for this variable; the remaining 10% represents the redundant effect of *patX* expression.

We consider the same functional mobile form of the inhibitor for all the inhibitory genes considered: *patS, hetN* and *patX*. This inhibitor is the hexapeptide ERGSGR [22]. We consider that the hexapeptide is produced as a modification of PatS, HetN or PatX at the cell membrane, with characteristic rates for each protein. The product of the modification is exported to the neighbor cell. This hexapeptide has shown to have a higher affinity with HetR than the pentapeptide originally proposed [15].

An important novel inclusion in the model we propose here is the requirement of a post-transcriptional transformation of HetR to act as a genetic regulator. This active form of HetR, which we term AHetR, is probably obtained through a phosphorylation process [29, 56, 58] and only the active fraction of HetR would contribute to heterocyst differentiation. This could explain the apparent paradox of a higher concentration of HetR with less heterocyst formation in both Δ*hetF* and Δ*patA*. The high concentration of HetR in these mutants would be explained through a higher turnover rate for the activated HetR protein [56]. A possible active form of HetR has been recently suggested in [59], where the authors present evidence for phosphorylation of HetR as crucial for its activity in *Nostoc PCC 7120*. This phosphorylation is shown to require the presence of the Pkn22 kinase but no more information regarding the regulation of this phosphorylation is provided. Here we hypothesize that this would be the role of the genetic pathway controlled by *hetF* which has already been presented as a protease [26] and therefore can be expected to have a role in post-transcriptional modifications. Thus, we assign to *hetF* the role of activator of HetR with the mediation of PatA through a post-transcriptional interaction that would change HetR into its enzymatic form. This hypothesis is inferred by observing changes in the phenotypes of several mutant backgrounds and has been considered in Fig 1. However, the nature of this interaction cannot be confirmed, since the observed phenotype could also be explained through indirect interactions mediated by HetF.

To complete the mechanistic model, one must take into account that different genes are expressed in heterocysts or vegetative cells as depicted in Fig 2. Despite the fact that it has also been observed in a tetramer configuration [58], there is still no experimental evidence that HetR binds DNA as a tetramer. Therefore, it will be assumed that it binds to the DNA only as a dimer [57,75]. Nevertheless, we have checked that this alteration does not change the qualitative system behavior and the same dynamics can be recovered assuming tetrameric binding with an alternative set of parameter values.

As shown in Fig 2, while *hetF* is produced only constitutively at a low-level [35], *patA* only has a regulated expression that depends on the active form of the dimeric transcription factor AHetR, which also activates its own expression. At the protein level: the HetR dimer needs to be activated by HetF (whose enzymatic activity can be enhanced by PatA) to become AHetR. PatS becomes an inhibitor of the transcription factor by protein transformation during cell to cell transport. The inhibitor thus produced is a small molecule that can move along the filament. *hetN* is expressed basally in heterocysts and becomes an inhibitor of the transcription factor, similar to the PatS product, by protein transformation during cell to cell transport. The fixed nitrogen products produced by the heterocyst can also move to act as an inhibitor of AHetR.

Using the regulatory logic in Fig 2 we have built a mathematical model of gene regulation on a growing filament of cells, as detailed in Materials and Methods.

### Study of the wild-type and the Δ*patS* and Δ*hetN* mutants

The statistical distribution of vegetative cell intervals between heterocysts may differ from one experiment to another, as one can notice comparing the results from different authors [12–14,18,24,34,54,76,77]. For consistency, to compare our results with the experimental data for the wild-type and both the Δ*patS* and the Δ*hetN* mutants, we consider the relative frequency of vegetative intervals of a given length presented in [54], which is the most recent dataset available and one of the most comprehensive.

As presented in the Materials and methods section, in order to simplify the description we have modeled both genes *patX, patS* using a single variable. Thus, a complete loss of function of this variable represents the experimental Δ*patX*Δ*patS* mutant. This double mutant induces considerably more heterocysts than the single Δ*patS* mutant, S2.Fig, as observed experimentally [20] (see movie S1.Movie). We have also simulated the Δ*patX* mutant (see movie S2.Movie), where the production rate is only 10% of the combined PatS+PatX variable. The phenotype observed (S2.Fig and S3.Fig) is still compatible with the wild-type data, as reported in [20], with both slightly shorter vegetative intervals and a higher percentage of heterocysts. In Fig 3 we observe the agreement between the simulations and the experimental data from [54] for wild-type and *patS* and *hetN* mutants.

**Fig 3.**
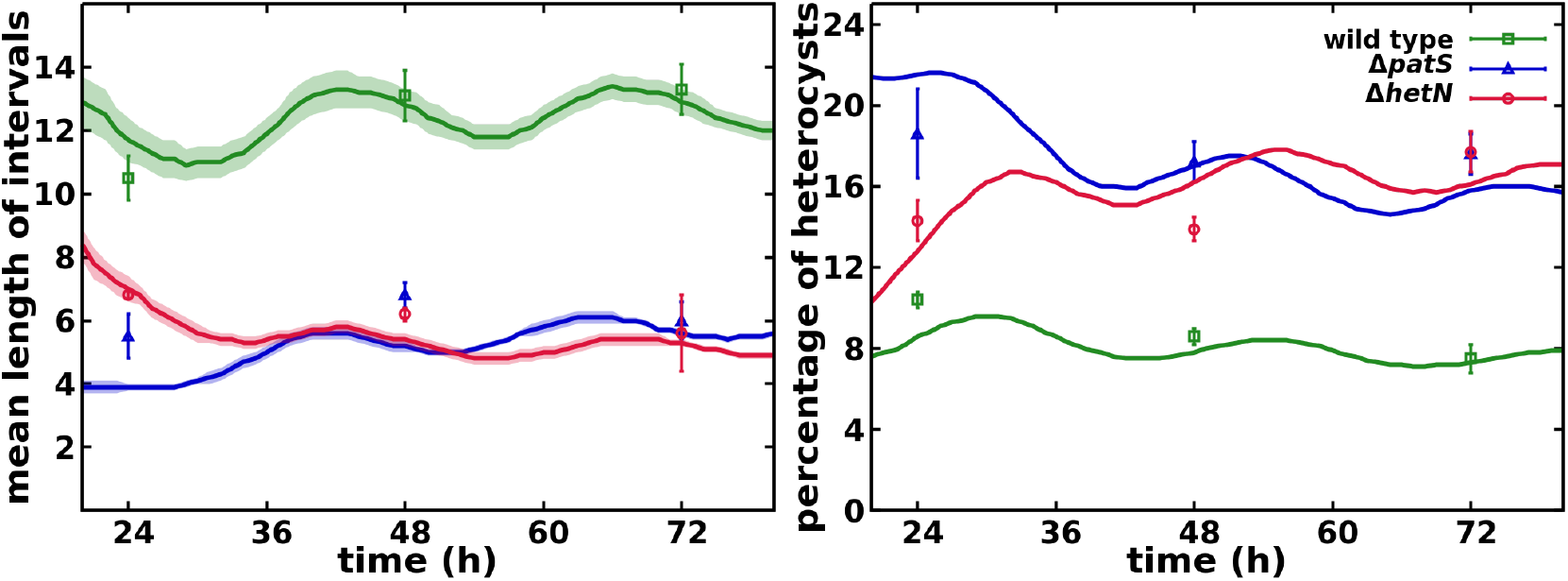
Mean number of vegetative cells between heterocysts and total percentage of heterocysts in the filament. Symbols represent experimental values from [54] and lines are simulation results.

For the wild-type, it is well known that roughly one of every ten cells differentiates with a slight increase of this interval length with time (see movie S3.Movie). In the Δ*hetN* mutant, the initial pattern is similar to the wild-type, except for the contiguous heterocysts appearing at a sequential pace (see movie S4.Movie). The Δ*patS* mutant shows a cluster of heterocysts, which appear simultaneously at short times (see movie S5.Movie). For longer times, the pattern of heterocysts is more similar to the wild-type, but with a higher incidence of contiguous heterocysts, in agreement with the experimental results reported in [14] with a larger statistical sample.

The comparison of the distributions of the number of vegetative cells between heterocysts in the experiment of Ref. [54] and the simulated filaments are shown in Fig 4. The agreement is very good. A small deviation appears for the early phenotype of the Δ*patS* mutant, especially at 24h. The simulations present less contiguous heterocysts and shorter intervals than in the experiments. This difference could be due to the effect of not considering a protoheterocyst phase. Without this phase, the differentiation of adjacent cells is strongly reduced because, once a cell differentiates, it immediately starts producing both HetN and nitrogen products, which inhibit differentiation. Thus, an artificial surplus of one and two cell intervals is observed in the first round of differentiation. After the first round of division, this causes the observed peaks of two and four cell intervals observed at 24h as observed Fig 4 and the movie S5.Movie.

**Fig 4.**
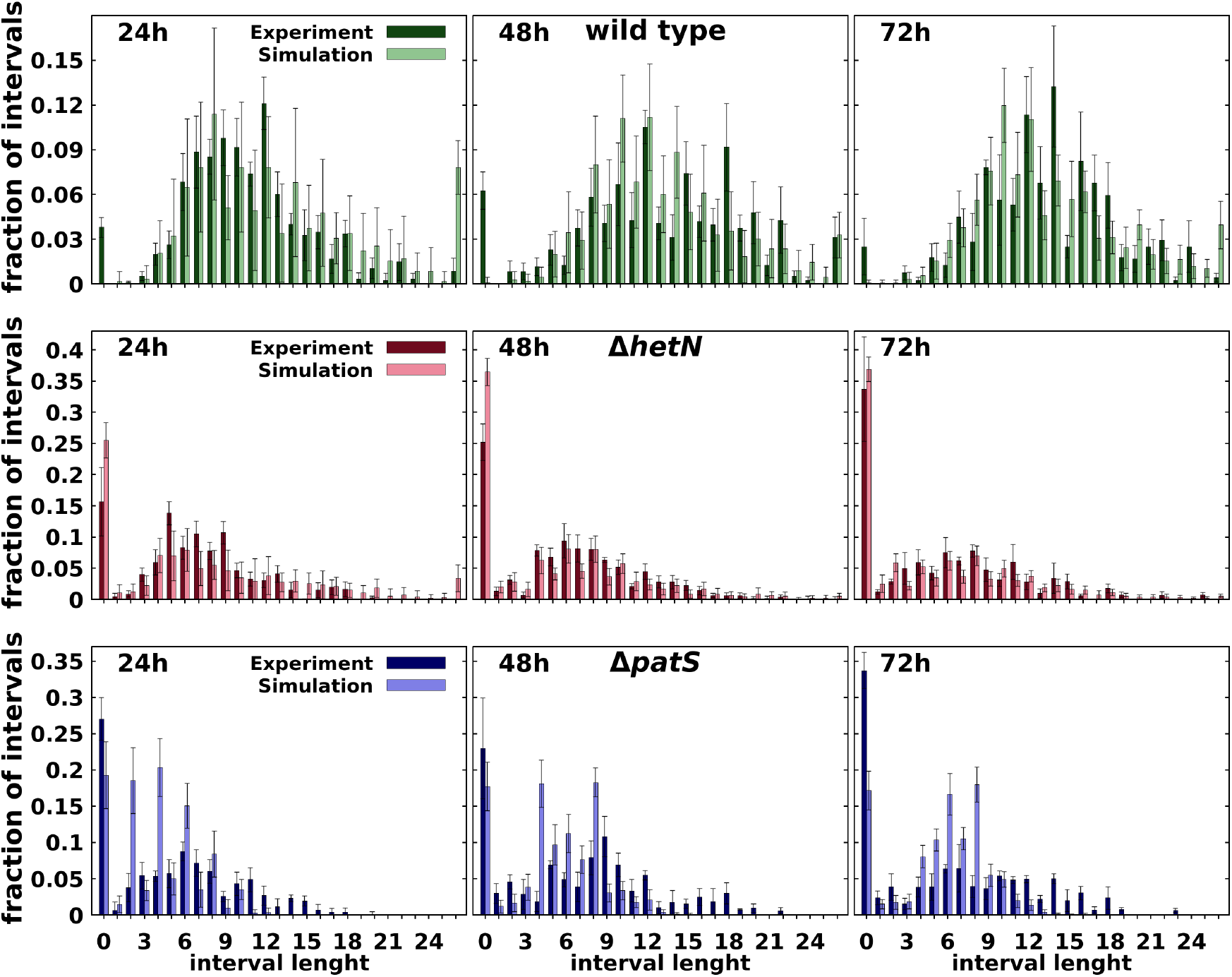
Comparison at different times after nitrogen deprivation (as indicated) between experimental [54] and simulated histograms of the number of vegetative cells between heterocysts for wild-type (top row), Δ*hetN* (middle row), Δ*patS* (bottom row).

The low amount of contiguous heterocysts observed in wild-type simulations in comparison with experimental data [54] can be explained using the same argument. A protoheterocyst phase would ease the stochastic formation of multiple simultaneous heterocysts in all genetic backgrounds. On top of that, it is worth noting that these contiguous heterocysts are seldom described in the literature. Their appearance on the wild-type can be explained by stochastic fluctuations of the genetic expression that get fixed through the irreversible process that is the differentiation into a heterocyst. In any case, the model also allows for the formation of these contiguous heterocysts, albeit in a much smaller proportion.

These results also improve the phenotypes obtained in the minimal model from [47]. In that model, the Δ*hetN* mutant showed both smaller clusters of heterocysts and shorter intervals between them. Additionally, this previous model could not produce any contiguous heterocysts in the wild-type.

### Loss of heterogeneity in the HetR concentration profile in a Δ*patA* mutant background

The simulations for the Δ*patA* single mutant show a similar behavior to experimental results [12, 26, 27, 30, 79], with heterocysts forming mostly on the filament ends despite higher global HetR concentration in the filament (approximately 24% higher in our simulations, Fig 5) in Δ*patA* than in the wild-type.

**Fig 5.**
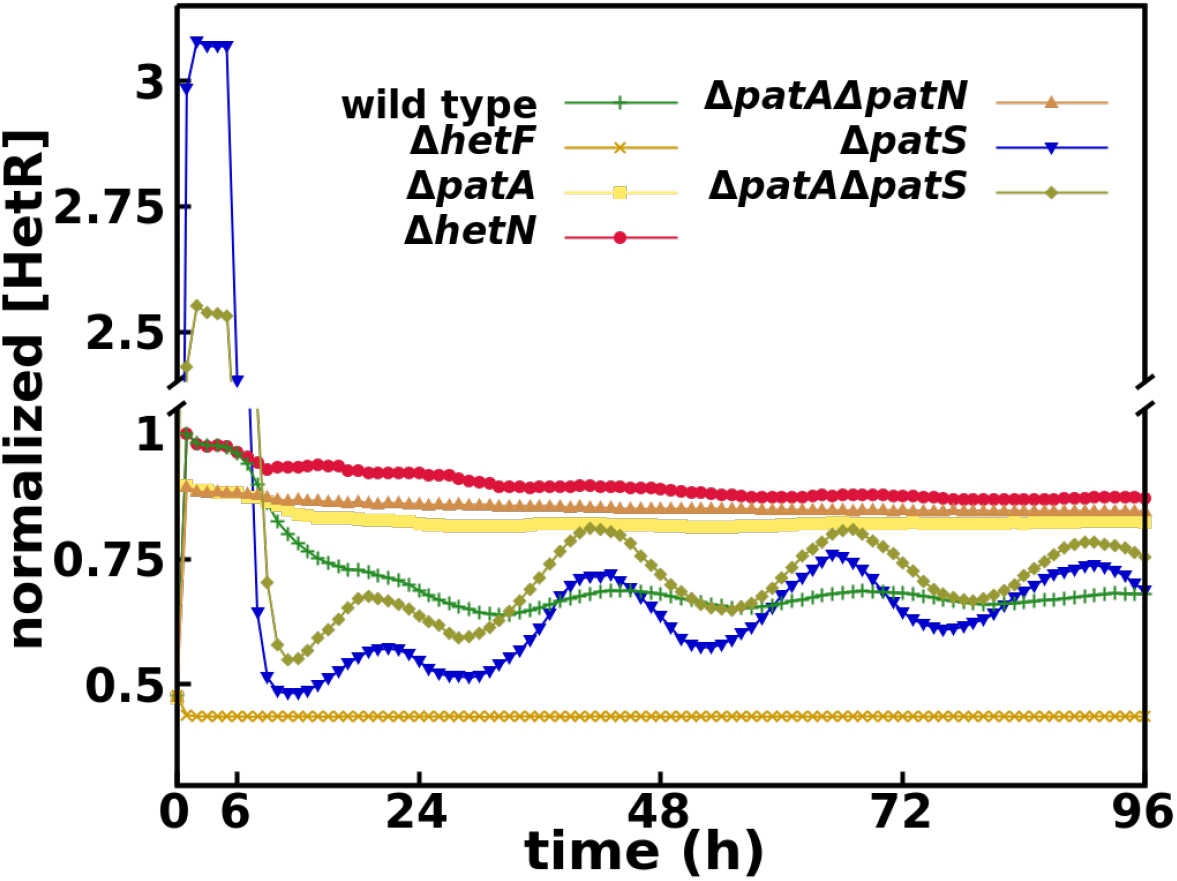
Time evolution of HetR concentration on vegetative cells for all mutants considered, normalized by the maximum wild-type concentration.

The absence of internal heterocysts is caused by the loss of a distinct HetR concentration profile, Fig 6 and movie S6.Movie. The absence of PatA in the system slows the conversion to active HetR to a minimum level and therefore also the regulatory effect of HetR over both itself and *patS*. This produces a homogenization of the production of PatS and also of the inhibitory hexapeptide along the filament. Then, in the absence of a pronounced inhibitory gradient, the levels of HetR increase uniformly to levels close to the threshold for differentiation (Fig 6). In this conditions, the selection of the few internal cells that will differentiate is exclusively due to stochastic fluctuations on the protein production of both HetF and PatS. On the other hand, the model assumes a certain passive diffusion of both the inhibitory hexapeptide and fixed nitrogen through the filament ends, which causes a local reduction of the inhibitory signals, allowing the differentiation of the boundary cells. If one does not allow the diffusion through the border cells heterocysts do not form in the filament ends as HetR does not accumulate enough on those cells, Fig 6. A simple analogy to a continuous reaction-diffusion system (see Materials and methods) explains how the properties of molecular trafficking at the filament ends, boundary conditions in the mathematical language, can lead to different profiles of heterocyst differentiation.

**Fig 6.**
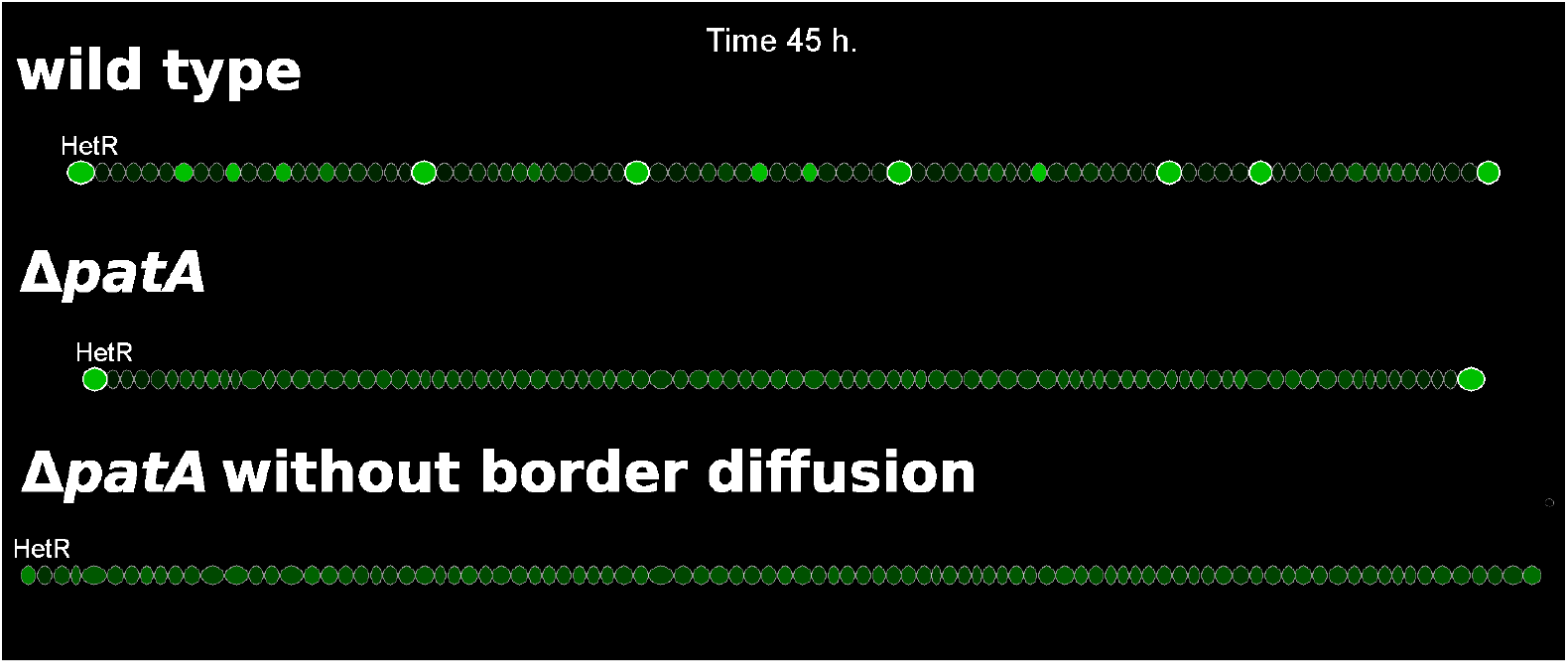
HetR profiles in filaments of wild-type, Δ*patA*, and Δ*patA* with no inhibitor leakage from the terminal cells.

The results of the simulation for the Δ*patA*Δ*hetN* double mutant also show a phenotypic agreement with observations [12]. The filaments present multiple terminal heterocysts with only the occasional internal heterocyst (movie S7.Movie). Additionally, the model predicts a higher HetR concentration (approximately 27% higher in our simulations, Fig 5) in Δ*patA*Δ*hetN* than in the wild-type, equivalent to the one observed in Δ*patA*. Here the border effect of the inhibitor diffusion through the ends of the filament gets propagated to multiple cells because the Δ*patA*Δ*hetN* double mutant, besides the homogenization of the HetR concentration, does not present the inhibitory gradient around existent heterocysts produced by *hetN*.

In Fig 7 we present histograms of the amount of terminal heterocyst for the Δ*patA*Δ*hetN* mutant. These results are in agreement with experimental data [12]. The model seems to have a small delay in the formation of the heterocysts (also present in the Δ*patA* mutant, data not shown). This delay could be related to the mechanism of commitment to the differentiation of a given cell. In our model, this decision is exclusively linked to a sustained high concentration of HetR and not to the expression of a supplementary gene (*hetP* and/or *hetZ*) as recent publications [60,61,81–84] seem to indicate. Hence, incorporating a gene-controlled differentiation commitment would surely improve these results, because then self-regulation of that gene could amplify the differentiation signal.

**Fig 7.**
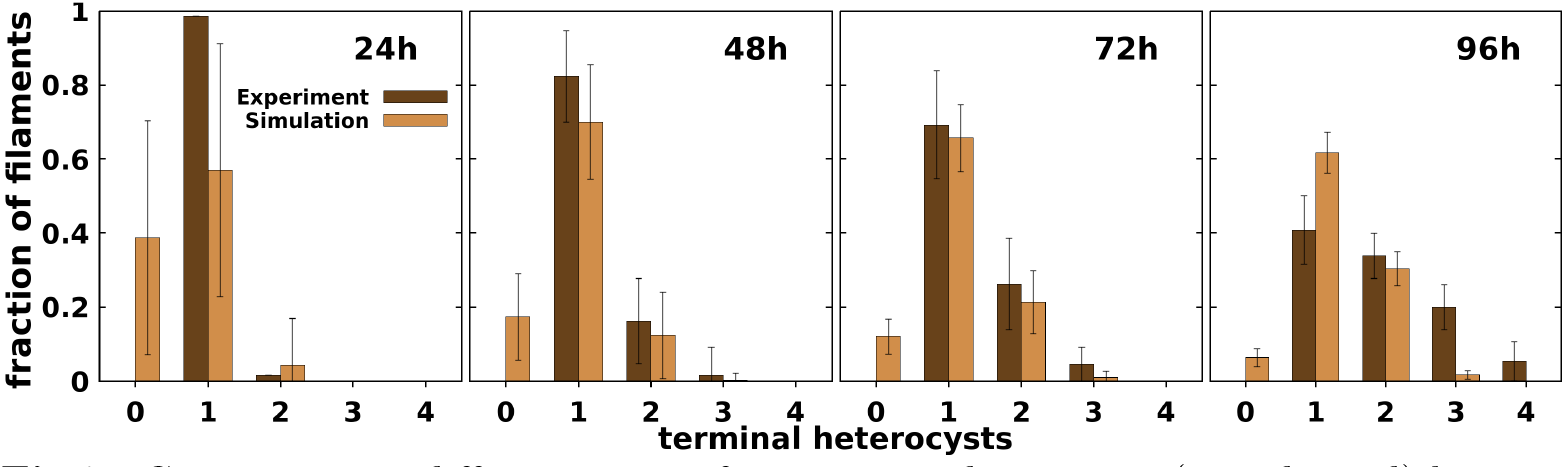
Comparison at different times after nitrogen deprivation (as indicated) between experimental [12] and simulated histograms of the number of heterocysts in the filaments ends for the Δ*patA*Δ*hetN* double mutant.

Finally, we studied the Δ*patA*Δ*patS* double mutant, of which to our knowledge there are only phenotypical observations [12]. Its phenotype is described as similar to the single Δ*patS* mutant phenotype, but with longer distances between heterocysts. The simulated filament (movie S8.Movie) fits this behavior, S2.Fig and S4.Fig. One can notice that the Δ*patA*Δ*patS* mutant presents a higher amount of contiguous heterocysts but with a smaller frequency and larger intervals of vegetative cells between heterocysts. Additionally, the simulated filament does not present an increase in HetR concentration typical to other *patA* mutants. This also confirms that the full Δ*patA* phenotype, which shows almost no internal heterocysts and a high concentration of HetR, requires a functional *patS* gene [12].

As we discussed previously, the increase of the HetR concentration is produced due to the homogenization of the PatS production along the filament, which produces a state where all the cells have a homogeneous HetR concentration lower than the decision threshold. Therefore if *patS* is not functional this effect would not be observed and the weak activation of HetR due to the absence of PatA precludes the formation of as many clusters of heterocysts as one observes in the Δ*patS* mutant.

One can observe this in Fig 5, where the HetR concentration for this mutant does not behave like the Δ*patA* single mutant or the Δ*patA*Δ*hetN* double mutant, with high constant values of HetR. It behaves similarly to the simple *patS* mutant, with a slightly higher concentration than that mutant after the first outburst of HetR that marks the first differentiation round [35]. On the other hand, the concentration is higher in the Δ*patS* single mutant during the first round of differentiation. This inversion is due to the faster production of HetR with a functional *patA*. Therefore, the decision to differentiate is reached in a shorter time than in the Δ*patA*Δ*patS* mutant. Hence, on a homogeneous initial condition (the first round of differentiation), the concentration would be higher in the Δ*patS* mutant because all cells start producing HetR at a much faster rate. But starting from heterogeneous initial conditions (all the following differentiation rounds), the concentration would be higher in the double mutant. Due to the slower commitment, more cells (closer to the heterocysts) start producing HetR before being shut down by the newly formed heterocysts. This effect can also be observed by comparing movies S5.Movie and S8.Movie.

### Complete loss of HetR regulatory function in the ΔhetF mutant background

If HetF is necessary to produce the active form of HetR, in its absence HetR should lose its regulatory function. This prediction is in agreement with experimental observations in *Nostoc punctiforme* [35], which presents a similar transcriptional induction pattern of *hetR* than *Anabaena* PCC 7120. There, the induction of *hetR* is dependent on the presence of an intact copy of *ntcA*. Moreover, the induction of *hetR* is still present in the Δ*hetF* background but with an altered induction pattern that eliminates the 0 to 6h burst.

Our simulated Δ*hetF* successfully eliminates the initial burst of HetR production observed in all other mutants, Fig 5. The absence of this burst in Δ*hetF* implies that it is exclusively produced by the positive self-regulation of *hetR* [26, 35]. Due to this, the introduction of an additional *hetR* promoter activated as a response to nitrogen deprivation would improve the agreement between our model and experiments by increasing only the overall HetR concentration in our simulations (especially on the Δ*hetF* and Δ*patA* simple and multiple mutants) without altering much of the dynamics.

## Conclusion

The formation and maintenance of heterocyst patterns is a paradigmatic example in which many processes, such as complex gene regulatory networks, interactions at different time scales, molecular trafficking, and cell growth, act together to give rise to a multicellular pattern. All these aspects form an intriguing puzzle for which a complete understanding is still elusive. A practical way to expand our knowledge about this problem is to investigate what are the functions of some of its pieces. Thus, increasing the complexity of a previous minimal model, we have been able to gain insight into the functions of the players involved. We have focused on the *patA* gene, and based on experimental phenotypical evidence, we have formulated a mathematical model that includes the interactions of this gene with the key genes responsible for heterocyst pattern formation. Our model shows that considering PatA as a collaborator of the activation process of HetR directed by *hetF* is capable of explaining all the phenotypes of the genes considered in our genetic network. This agreement suggests that there is some interaction, direct or indirect, between *hetF* and *patA* that has not been reported experimentally. This consideration, together with the existence of an active form of HetR, is also enough to account for the paradoxical changes in HetR concentration in the *patA* mutant, that seemed to question the central role of *hetR* in heterocyst differentiation.

New experiments are required to confirm the validity of the interactions proposed. In particular, experimental information regarding protein translation [25, 69] could be useful in order to have more detailed information regarding the effects of a given gene on the regulatory network. The roles of many other actors are still to be elucidated and could be included in the core processes to obtain a more extensive mathematical description. For example, recent work [72] presents evidence that *hetL*, a gene previously shown to alter PatS mediated inhibition of heterocyst differentiation [73], interacts with HetR without inhibiting its DNA-binding activity in *Nostoc PCC 7120*. This interaction protects HetR from the inhibitory effects of the Pat-derived hexapeptide and seems to be essential for the proper function of HetR as a genetic regulator. That would be the role that we have assigned to hetF based on the phenotypic evidence on *Anabaena PCC 7120*.

Finally, our work shows that it is possible to reproduce the *patA* mutant phenotype without considering a differentiation mechanism depending on a cell’s position on the filament. The model also expands the characterization presented in [12] by directly linking the increase of the HetR concentration in all the cells with the absence of internal heterocysts in both Δ*patA* and Δ*patA*Δ*hetN* mutants. This is obtained by a slowing of the *hetR* transcription which produces a homogenization of HetR concentration through the filament. Then it is easy to see why this phenotype is not present in the Δ*patA*Δ*patS* mutant where this reduction on the transcription rate is completely compensated by the absence of the early inhibitor PatA. The intriguing differentiation of only terminal heterocysts appears as a consequence of the boundary conditions of the system: leakage of inhibitors out of the filament through the terminal cells. Hence, despite the apparent simplicity of *Anabaena* compared to other developmental systems, it is already clear that genetic and metabolic interactions result in patterns shaped by physical constraints.

## Supporting information

**S1.Table.** Model parameter values table. Parameter values used for the wild-type simulations. The abbreviation DL is used for dimensionless variables.

| Parameter | Description | Value | Units |
| --- | --- | --- | --- |
| $\beta_R$ | Basal production rate of HetR monomers | 66.665 | $nM \cdot h^{-1}$ |
| $\rho_R$ | Maximum regulated production rates of HetR monomers | 763.33 | $nM \cdot h^{-1}$ |
| $\alpha_R$ | Linear degradation rate of HetR monomers | 2.1293 | $h^{-1}$ |
| $\mu$ | Nonlinear degradation rate of HetR dimers | $1.6354 \cdot 10^{-3}$ | $nM^{-1}$ |
| $\rho_A$ | Maximum regulated production rates of PatA | 3862.3 | $nM \cdot h^{-1}$ |
| $\alpha_A$ | Linear degradation rate of PatA | 3.823 | $h^{-1}$ |
| $\tau_A$ | Equilibrium constant of the PatA enhancement of the activation proces of HetR dimers | 81.132 | $nM$ |
| $F_R$ | Fraction of HetF activated HetR dimers | $1.2578 \cdot 10^{-4}$ | DL |
| $\rho_S$ | Maximum regulated production rates of PatS | 337.5 | $nM \cdot h^{-1}$ |
| $\alpha_S$ | Linear degradation rate of PatS | 2.0827 | $h^{-1}$ |
| $c_S$ | Rate of conversion of PatS to the ERGSGR hexapeptide through cellular diffusion | 8.1485 | $h^{-1}$ |
| $\rho_N$ | Maximum regulated production rates of HetN | 527.2 | $nM \cdot h^{-1}$ |
| $\alpha_N$ | Linear degradation rate of HetN | 2.6204 | $h^{-1}$ |
| $c_N$ | Rate of conversion of hetN to the ERGSGR hexapeptide through cellular diffusion | 5.7671 | $h^{-1}$ |
| $\alpha_I$ | Linear degradation rate of ERGSGR hexapeptide | 2.3684 | $h^{-1}$ |
| $d_I$ | Diffusion rate of the ERGSGR hexapeptide | 9.0777 | $h^{-1}$ |
| $\rho_G$ | Maximum regulated fixation rates of nitrogen | 7533.7 | $nM \cdot h^{-1}$ |
| $\alpha_G$ | Linear degradation rate of fixed nitrogen | 4.32 | $h^{-1}$ |
| $d_G$ | Diffusion rate of the fixed nitrogen | 100.04 | $h^{-1}$ |
| $K_G$ | Equilibrium constant for the inhibitory reaction between HetR and the fixed nitrogen | 162.03 | $nM$ |
| $d_{border}$ | Rate of diffusion through the border cells | 0.381 | DL |
| $\Omega_\Lambda$ | Noise strength in cellular growth and size | 0.081 | DL |
| $\Omega_\Phi$ | Noise strength in gene expression and initial concentration | 0.21 | DL |
| $K_d$ | Equilibrium constant for the inhibitory reaction between HetR and the ERGSGR hexapeptide | 7.36 | $nM$ |
| $\lambda$ | Cellular growth rate | 0.08 | $\mu m \cdot h^{-1}$ |
| $M_\Lambda$ | Average maximum cell size | 4 | $\mu m$ |
| $T_R$ | Average minimum HetR concentration threshold to diferenciate | 110 | $nM$ |
| $M_R$ | Average accumulated HetR concentration required to form an heterocyst | 1320 | $nM$ |
| $T_{min}$ | Minimum time to differentiate | 5 | h |

**S1.Fig.**
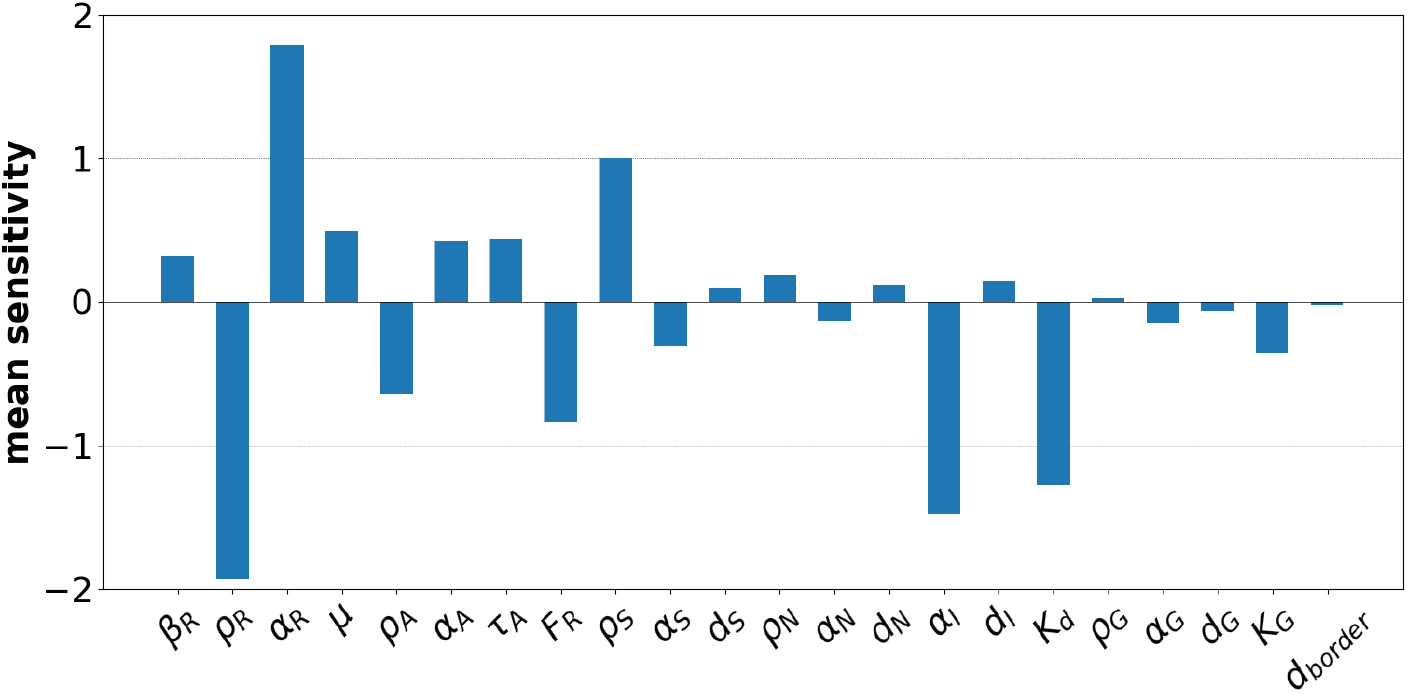
Sensitivity of the mean distance between heterocysts after 72h with respect to 10% changes in the indicated parameters. Changes are with respect to the wild-type 568 values in Table.S1.

**S2.Fig.**
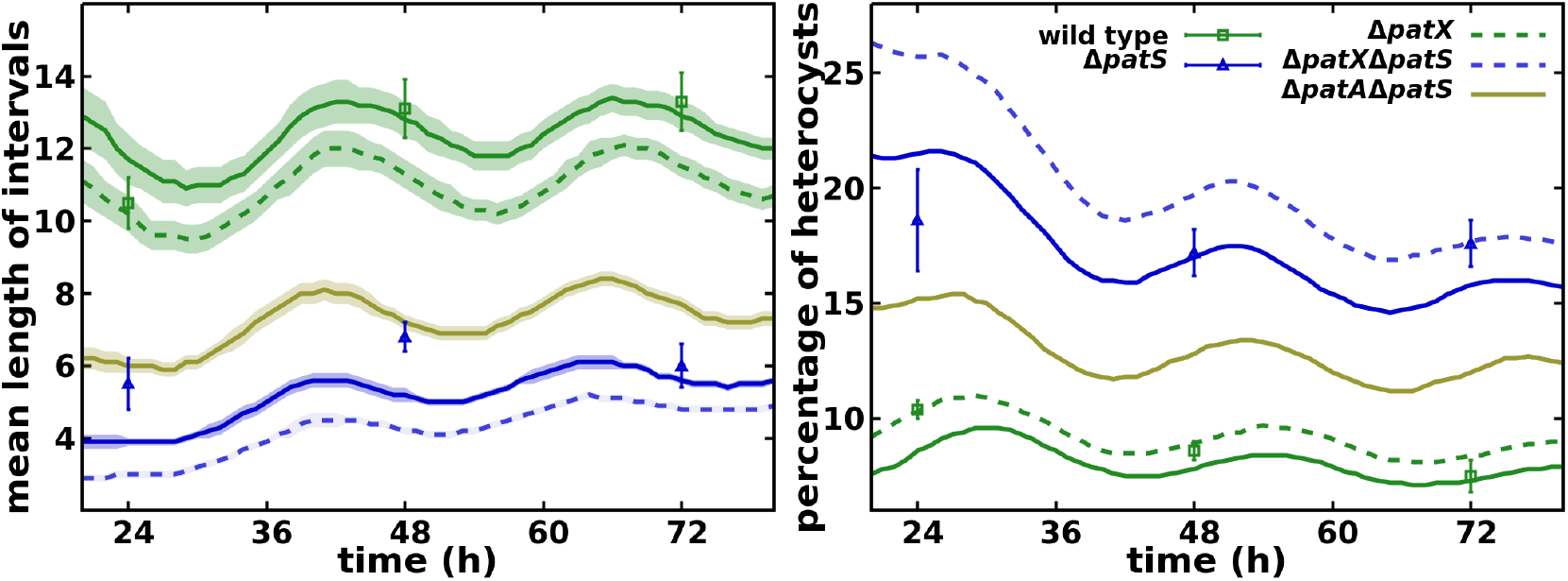
Mean number of vegetative cells between heterocysts and total percentage of heterocysts in the filament for different conditions, as indicated. Symbols represent experimental values from [54] and lines are simulation results.

**S3.Fig.**
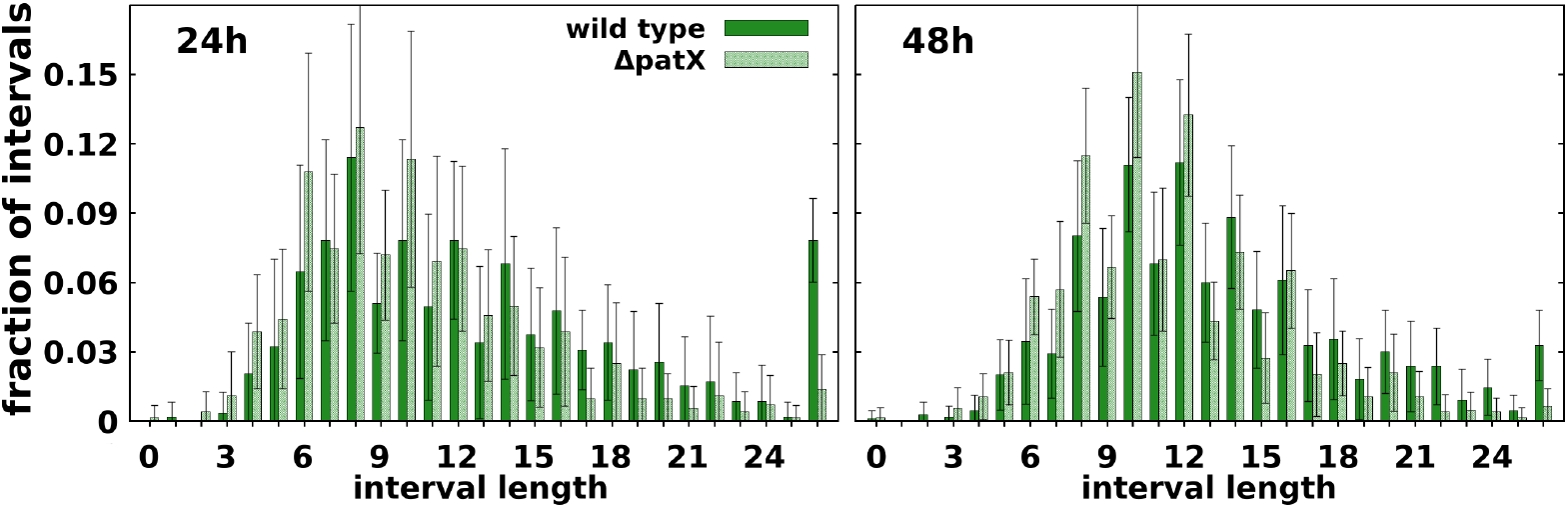
Comparison at different times after nitrogen deprivation (as indicated) between simulated histograms of the number of vegetative cells between heterocysts for wild-type and Δ*patX*, as indicated.

**S4.Fig.**
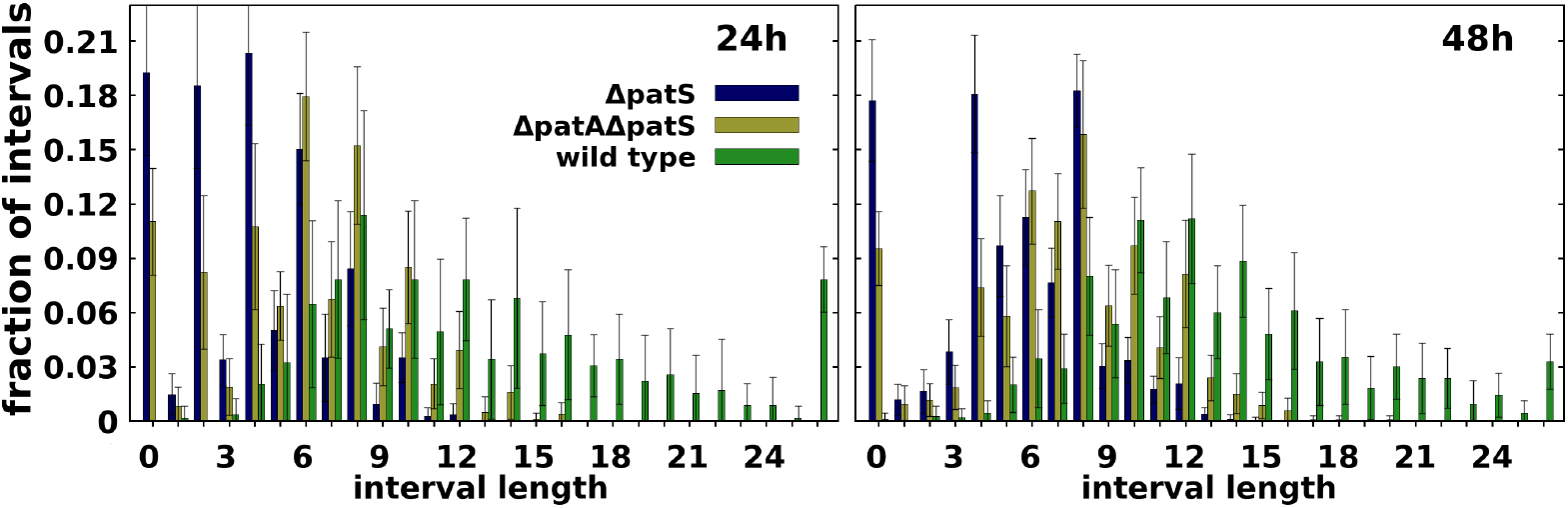
Comparison at different times after nitrogen deprivation (as indicated) between simulated histograms of the number of vegetative cells between heterocysts for wild-type, Δ*patS* and Δ*patA*Δ*patS*, as indicated.

**S1. Movie. Δ*patS*Δ*patX* simulation**

**S2. Movie. ΔpatX simulation**

**S3. Movie. wild-type simulation**

**S4. Movie. ΔhetN simulation**

**S5. Movie. ΔpatS simulation**

**S6. Movie. ΔpatA simulation**

**S7. Movie. ΔpatAΔhetN simulation**

**S8. Movie. ΔpatAΔpatS simulation**

## Acknowledgments

We thank Joel Stavans and Rinat Goren for fruitful discussions. This research was supported by MCIN/AEI/10.13039/501100011033/ and FEDER *Una manera de hacer Europa* through grant no. FIS2016-78313-P to S.A and the associated FPI contract BES-2017-079755 to P.C-F., and MCIN/AEI/10.13039/501100011033 through grant BADS, no. PID2019-109320GB-100, to S.A. and J.M-G. The Spanish MICINN has also funded the “Severo Ochoa” Centers of Excellence to CNB, SEV 2017-0712.

