## Supplementary figures and images for "Terminal heterocyst differentiation in the *Anabaena patA* mutant as a result of post-transcriptional modifications and molecular leakage"

### Supplementary Movie 1

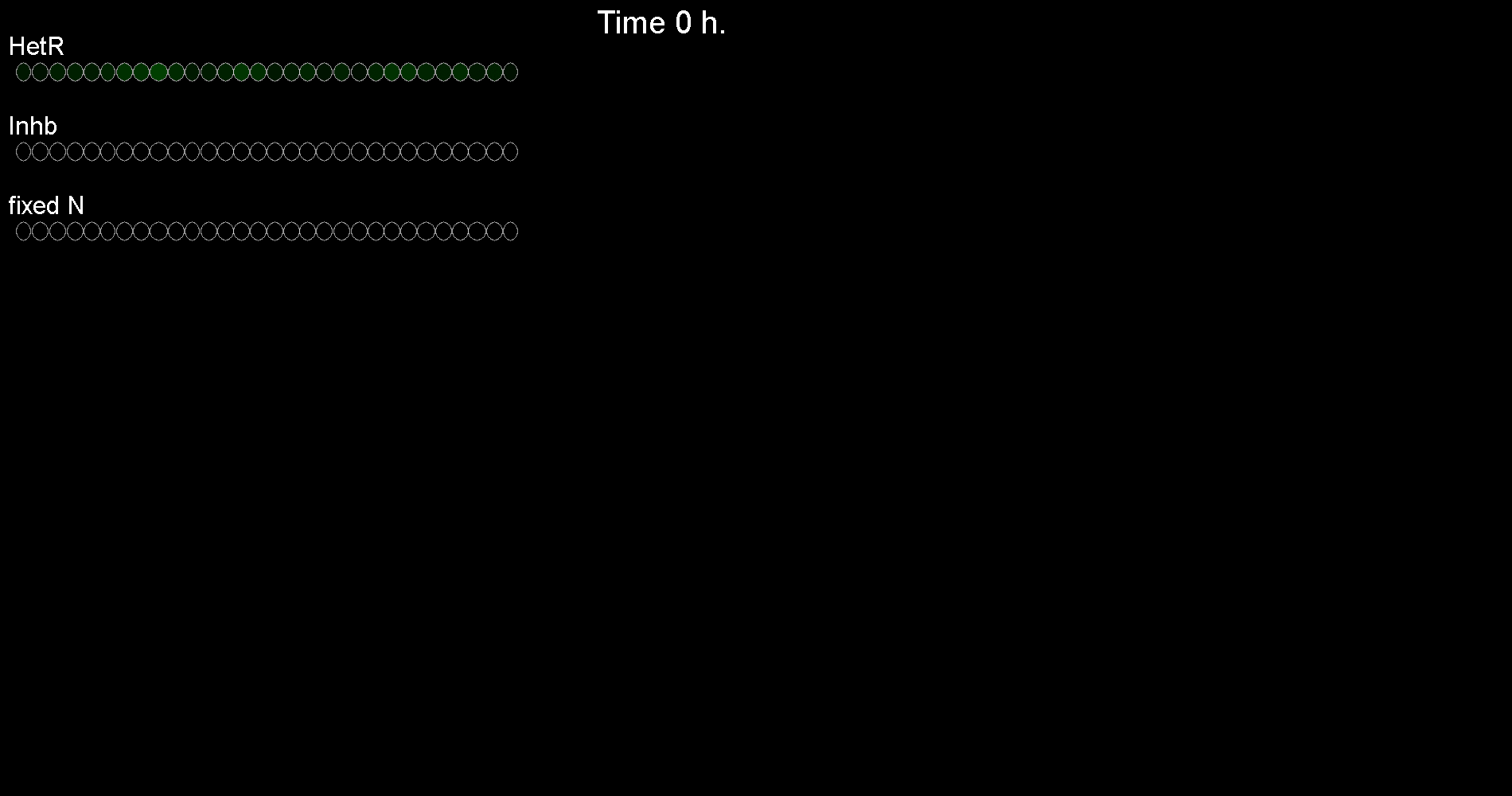

### Supplementary Movie 2

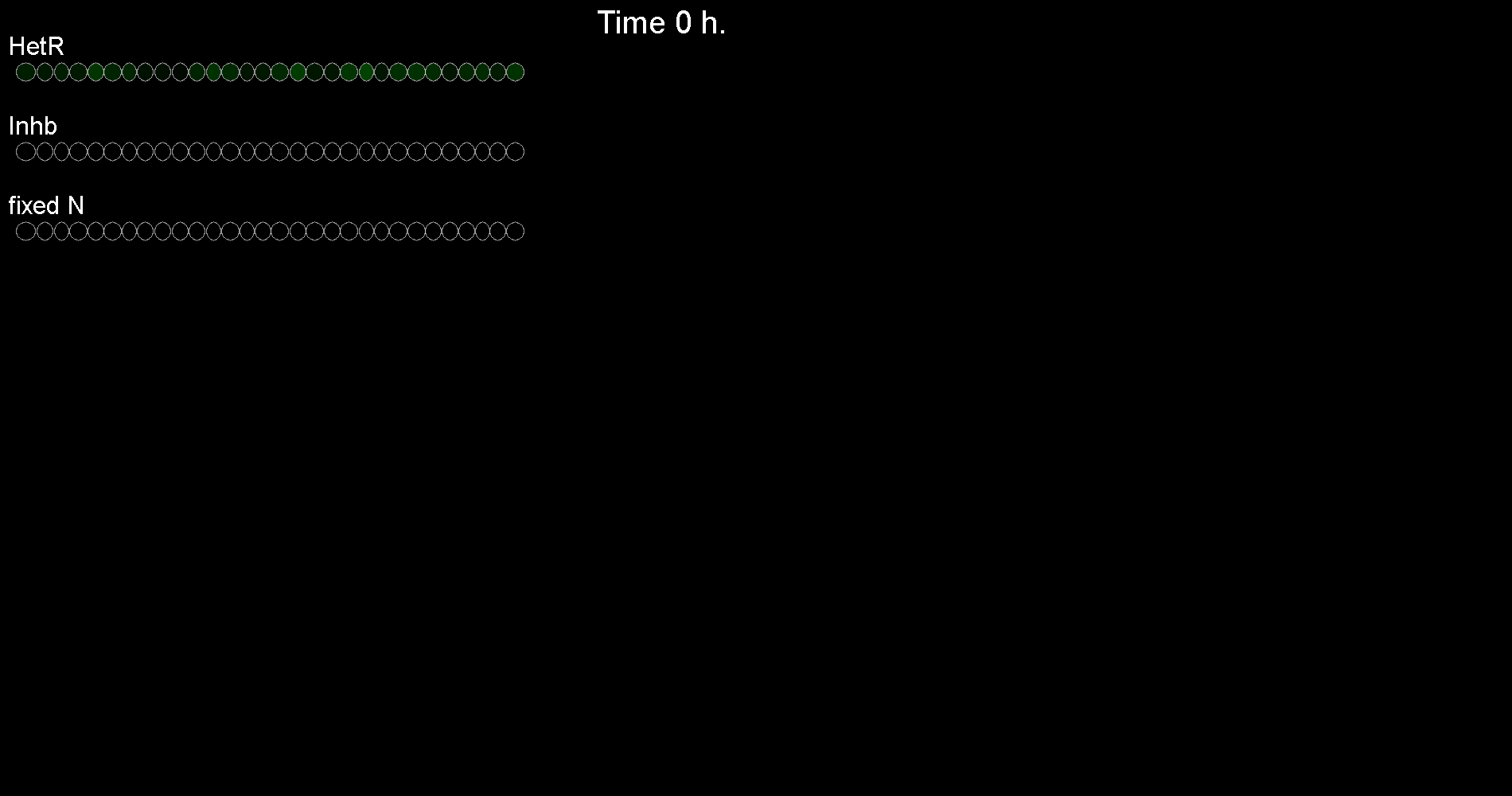

### Supplementary Movie 3

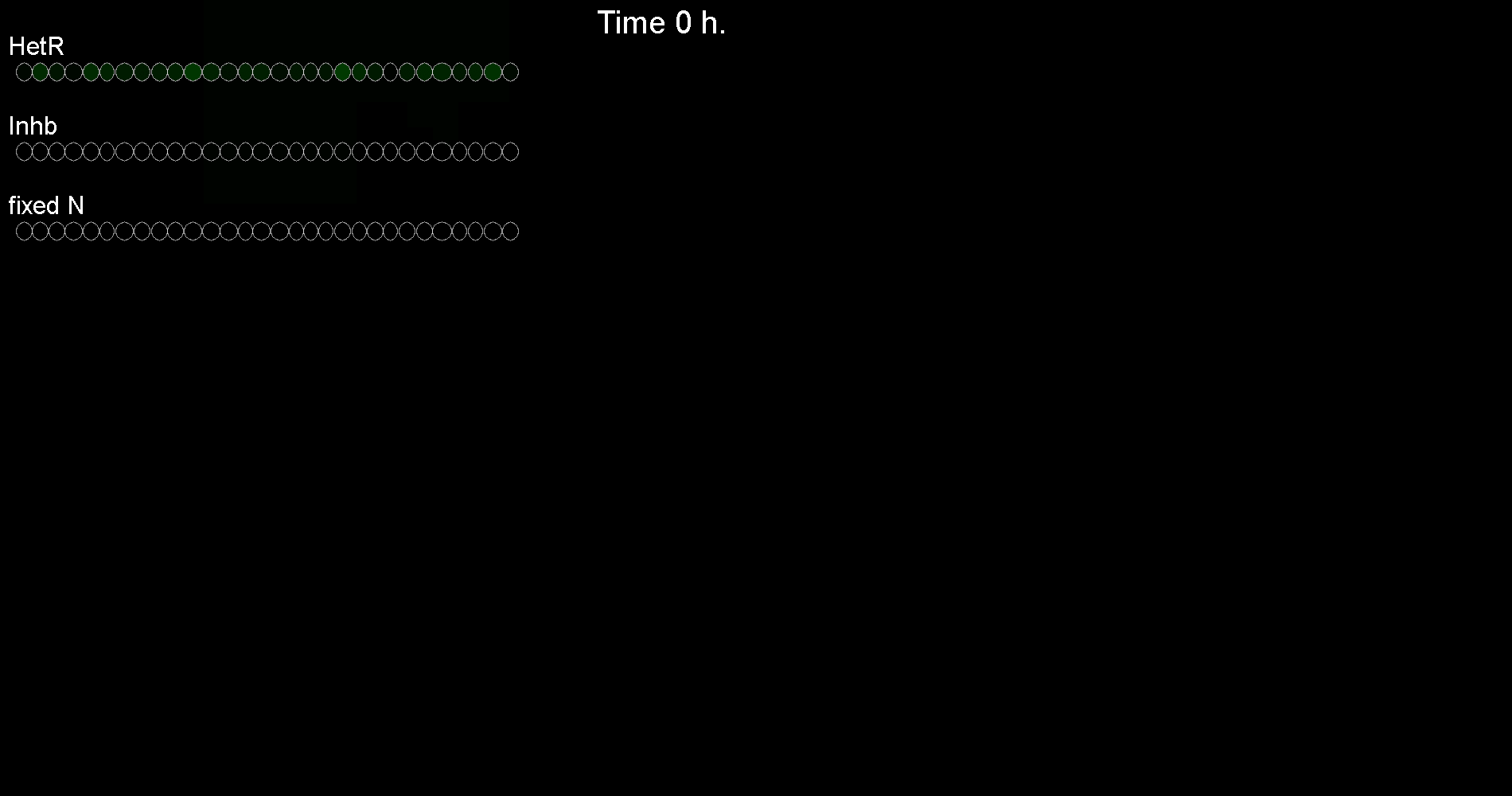

### Supplementary Movie 4

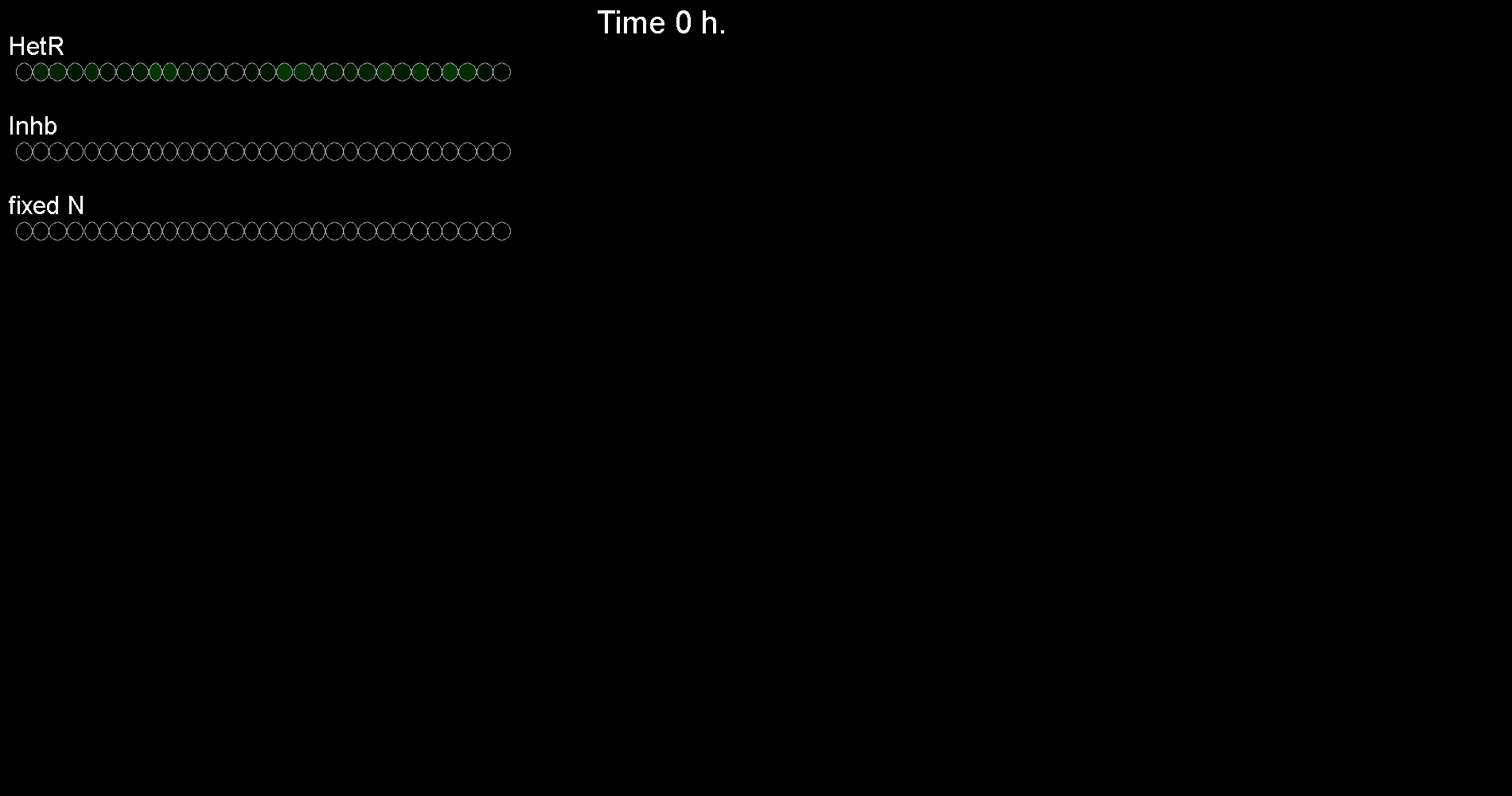

### Supplementary Movie 5

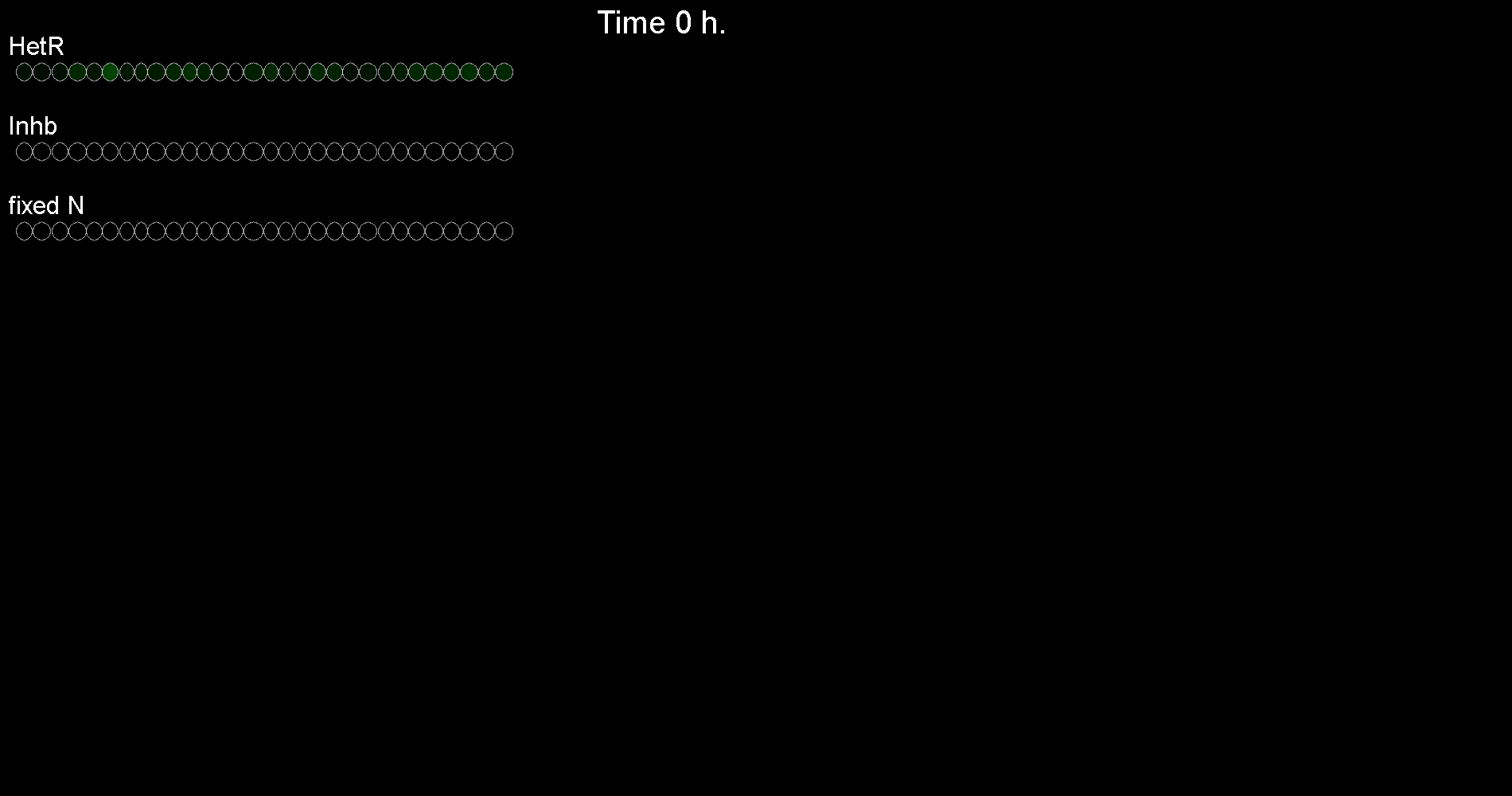

### Supplementary Movie 6

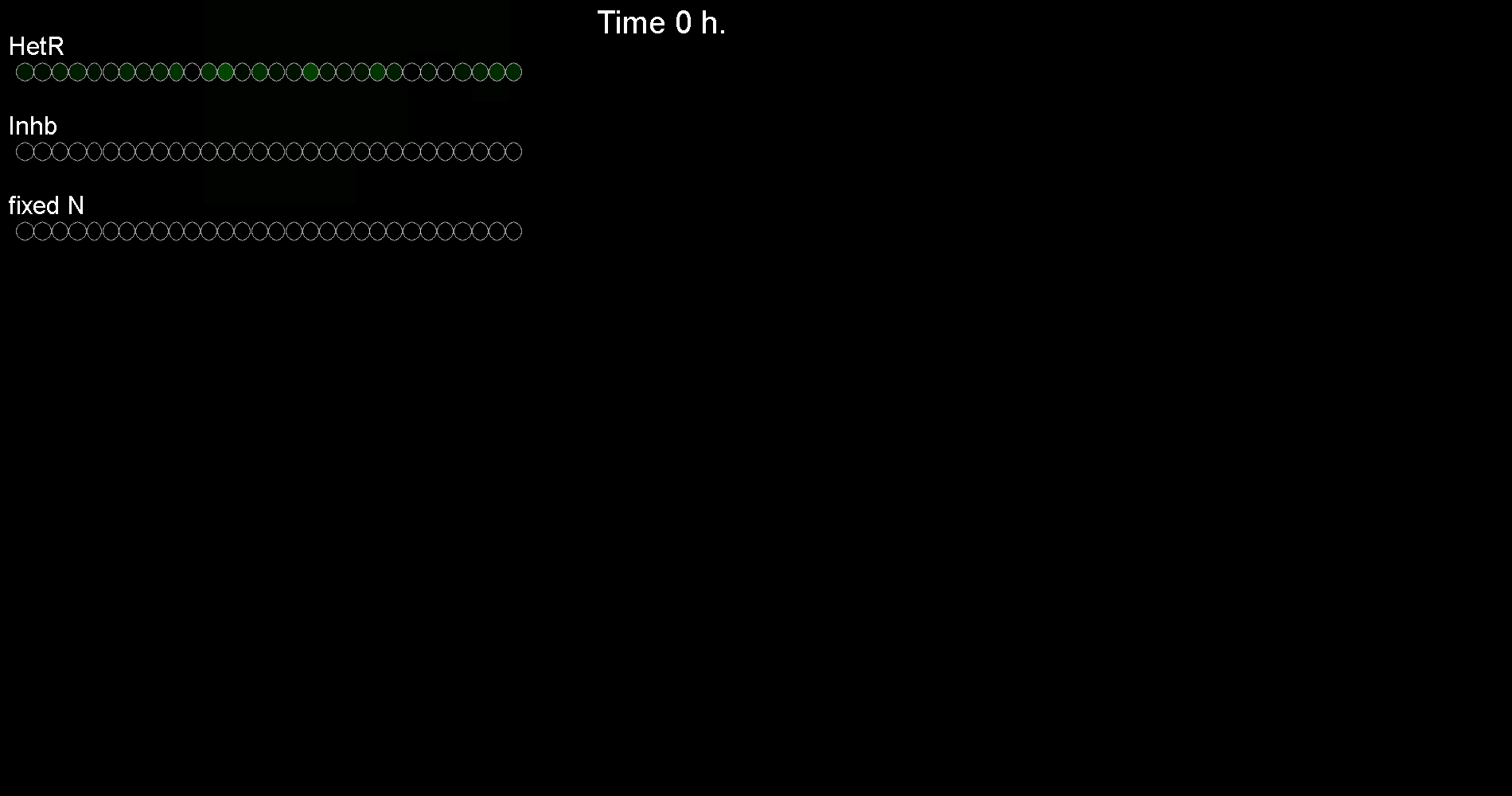

### Supplementary Movie 7

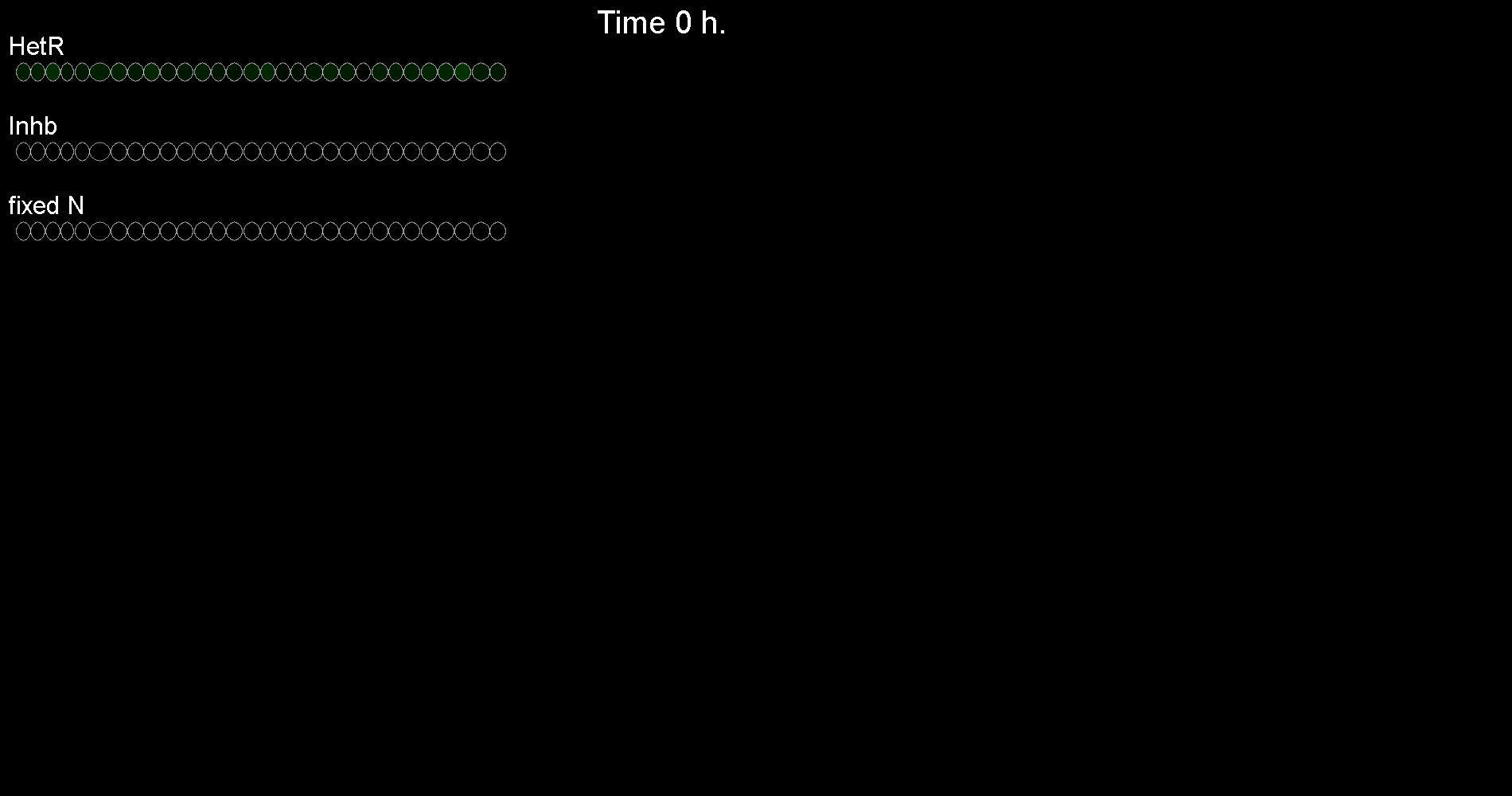

### Supplementary Movie 8

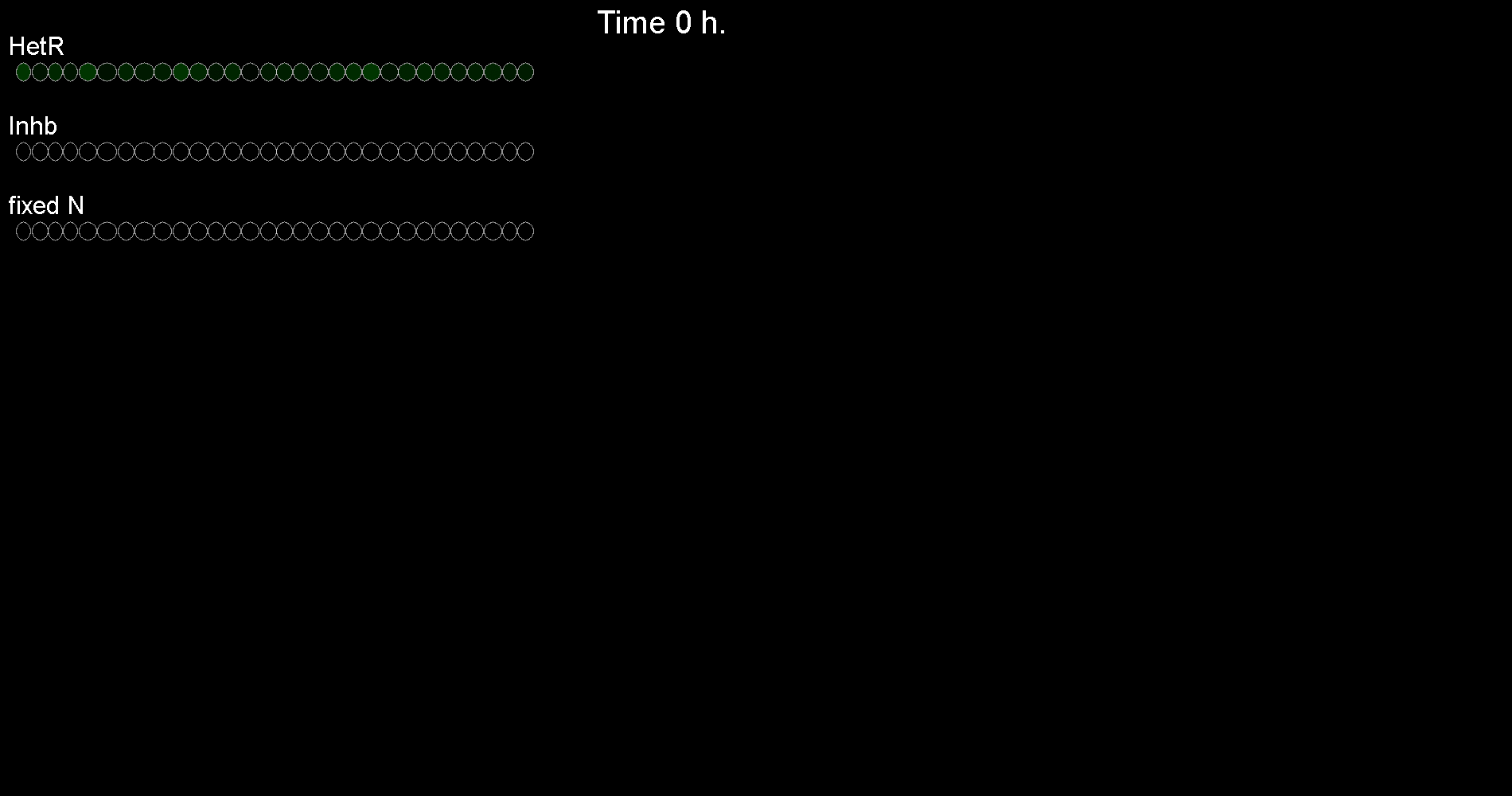
